# Structure of the Wnt-Frizzled-LRP6 initiation complex reveals the basis for co-receptor discrimination

**DOI:** 10.1101/2022.10.21.513178

**Authors:** Naotaka Tsutsumi, Sunhee Hwang, Simon Hansen, Deepa Waghray, Nan Wang, Yi Miao, Caleb R. Glassman, Nathanael A. Caveney, Kevin M. Jude, Claudia Y. Janda, Rami N. Hannoush, K. Christopher Garcia

**Author notes:** Equal contributions (Naotaka Tsutsumi, Sunhee Hwang, Simon Hansen).

## Abstract

Wnt morphogens are critical for embryonic development and tissue regeneration. Canonical Wnts form ternary receptor complexes composed of tissue-specific Frizzled receptors together with the shared LRP5/6 co-receptors to initiate β-catenin signaling. The structure of a ternary complex of an affinity-matured XWnt8-Frizzled8-LRP6 complex elucidates the basis of co-receptor discrimination by canonical Wnts by means of their N-termini and linker domains that engage the LRP6 E1E2 domain funnels. Chimeric Wnts bearing modular linker ‘grafts’ were able to transfer LRP6 domain specificity between different Wnts and enable non-canonical Wnt5a to signal through the canonical pathway. Synthetic peptides comprising the linker domain serve as Wnt-specific antagonists. The structure of the ternary complex provides a topological blueprint for the orientation and proximity of Frizzled and LRP6 within the Wnt cell surface signalosome.

## Introduction

Wnt morphogens play essential roles in embryonic development and tissue regeneration by orchestrating stem cell proliferation and differentiation (Clevers and Nusse, 2012; MacDonald et al., 2009; Nusse and Clevers, 2017; Rim et al., 2022). To initiate signaling, Wnt simultaneously engages the extracellular cysteine-rich domain (CRD) of a seven-transmembrane receptor Frizzled (Fzd) and an extracellular domain (ECD) of a single-transmembrane co-receptor LRP5 or LRP6, placing the two receptors in orientation and proximity conducive to activate downstream signaling pathways. The activated Wnt-Fzd-LRP5/6 complex inhibits the destruction complex, leading to the accumulation of the transcriptional co-activator β-catenin in the cytosol (MacDonald and He, 2012). The accumulated β-catenin translocates to the nucleus and induces the expression of Wnt target genes in concert with the T-cell factor/lymphoid enhancer factor (TCF/LEF) family of transcription factors.

Due to its crucial function in cell fate decisions, aberrant Wnt activation is linked to various diseases, including multiple types of cancers, making it an attractive therapeutic target by Wnt signaling antagonists (Anastas and Moon, 2013). On the other hand, Wnt agonism can be harnessed for regenerative medicine upon tissue injury or in bone and hair losses (Clevers et al., 2014). Engineered bispecific dimerizers that induce Fzd-LRP6 proximity can act as surrogate agonists in order to therapeutically exploit Wnt’s powerful functions in stem cell expansion and tissue renewal (Chen et al., 2020; Janda et al., 2017; Miao et al., 2020; Tao et al., 2019). These molecules, as well as Norrin (Chang et al., 2015; Xu et al., 2004), demonstrate the induced proximity between Fzd and LRP5/6 is the principal mechanism of signal activation. However, the signaling strength of such agonists appears highly dependent on the relative orientation and proximity of the Fzd-LRP6 heterodimer, and structure-activity relationships are not fully understood due to the absence of the natural Wnt-Fzd-LRP6 ternary complex.

The previous Wnt-Fzd crystal structures revealed the binding mode of canonical Wnts, *Xenopus* Wnt8 (XWnt8) and Wnt3a to their primary receptor Fzd, in which the hand-shaped Wnt pinches the Fzd CRD (Fzd_CRD_) with “thumb and index” fingers (Hirai et al., 2019; Janda et al., 2012). However, these structures fell short of recapitulating the fully competent cell surface signaling complex because they lack LRP5/6 that interact with Wnt through their ECD consisting of four YWTD β-propellers flanked by EGF-like domains. Furthermore, it has been shown that Wnts can be clustered into different families based on their binding preferences to distinct LRP6 domains (Bourhis et al., 2010; Gong et al., 2010). It remains to be determined how different Wnts, which are highly conserved can bind to different regions in LRP6 in contrast to the fact that Wnts bind to the Fzd_CRD_ in a very conserved manner. Furthermore, it is also unclear how non-canonical Wnts have structurally evolved to bypass LRP5/6 co-receptor requirements for their function. For these reasons, structural information on a Wnt-Fzd-LRP6 ternary signaling complex is important for understanding the geometric requirements (*i*.*e*., orientation and proximity) of the receptors for signaling initiation, how Wnts differentially bind to LRP6, and finally, for rationally optimizing the surrogate Wnts to achieve the natural topology of the endogenous Wnt signaling complex.

## Results

### Wnt engineering

To reconstitute a stable Wnt-Fzd-LRP6 complex for cryo-electron microscopy (cryo-EM) analysis, we first sought to engineer a Wnt that binds with increased affinity to LRP6. However, since Wnt proteins are lipidated and insoluble in the absence of detergents and also require the dedicated cargo receptor Wntless (WLS) for secretion (Bänziger et al., 2006; Nile and Hannoush, 2016; Nygaard et al., 2021; Zhong et al., 2021), they are problematic for conventional cell surface display methods commonly used for soluble proteins (Janda et al., 2017). Thus, we devised a strategy to display a library of *Xenopus* Wnt8 (XWnt8) variants bound to human Fzd5_CRD_ (hFzd5_CRD_) on the cell surface of mammalian cells and select XWnt8 variants that bind with higher affinity to the ECD of LRP6 (Figure 1A top panel). In order to prevent Wnt-induced downstream signaling and internalization (Figure 1A top left panel), we replaced the transmembrane region of hFzd5 with a single transmembrane helix (Chesnut et al., 1996) (Figure 1A top right panel), which improved the cell surface display of XWnt8 (Figure 1A bottom panel). The cell-surface XWnt8-hFzd5_CRD_ complex robustly bound human LRP6 E1E2 (hLRP6_E1E2_), the Wnt8 binding module (Gong et al., 2010; Ren et al., 2021), which was tetramerized to increase the affinity through avidity effect for detecting by flow cytometry (Figure 1B). Deleting the NC-linker of XWnt8 (XWnt8_ΔNC_), which has previously been reported to mediate LRP6 binding in the context of Wnt3 signaling (Chu et al., 2013; Hirai et al., 2019), substantially decreased, but did not completely abolish, hLRP6_E1E2_ binding (Figure 1B). These indicated that the wild-type XWnt8-Fzd5_CRD_ complex displayed on the cell surface is functional regarding LRP6 recognition, and that the NC-linker is a primary LRP6 binding site, but additional interactions beyond the NC-linker may contribute to LRP6 binding. We then created ‘soft-randomized’ libraries of XWnt8 variants that harbored mutations in the NC-linker (residues 222-234) with an experimental diversity of approximately 1 million unique sequences (Figure 1—figure supplement 1A). The libraries were designed to introduce sparse mutations within the NC-linker that would exhibit enhanced binding energetics to LRP6 while maintaining the overall structural integrity of the linker.

**Figure 1.**
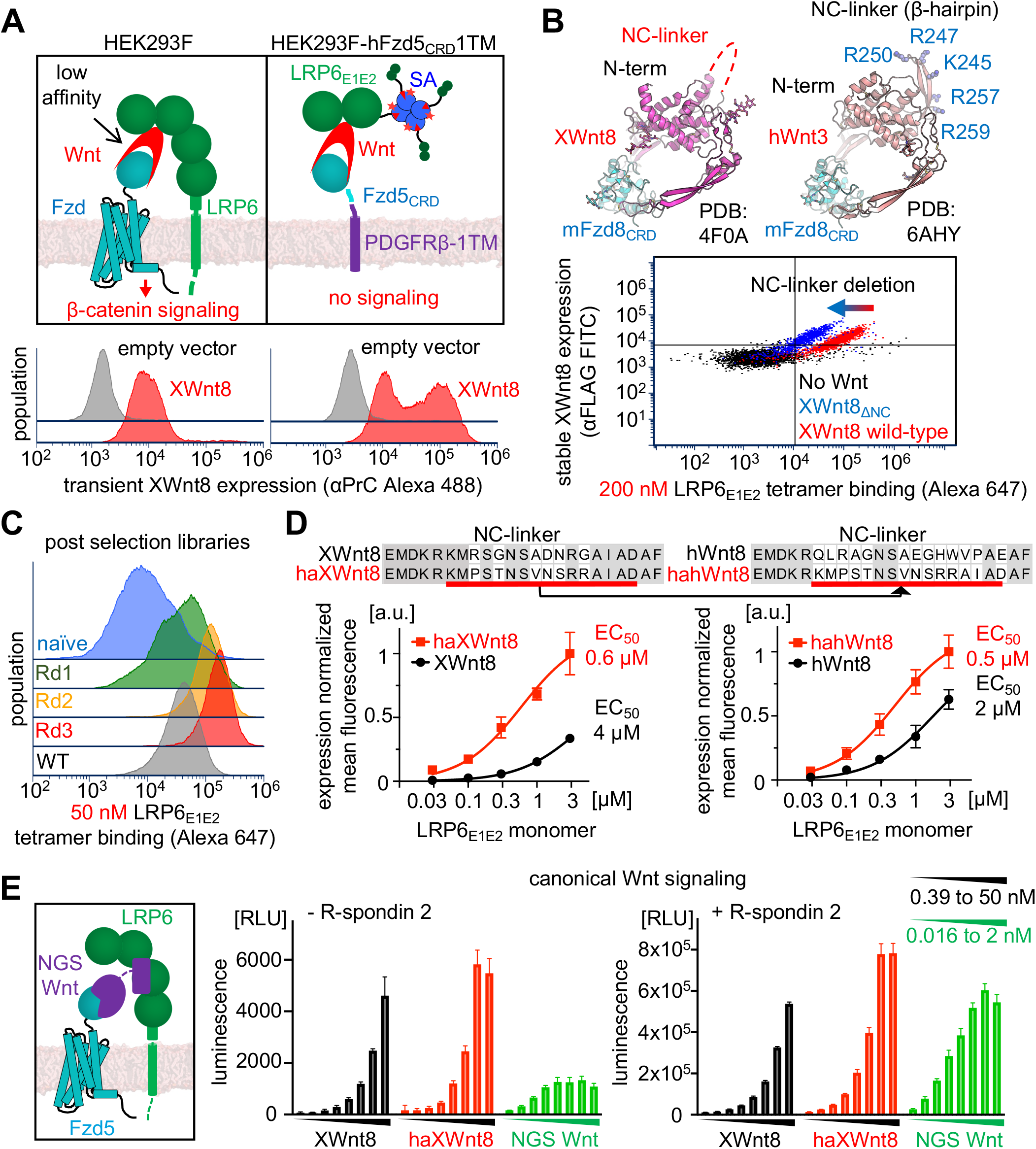
Engineering and characterization of haXWnt8. (A) Schematic for the cell-surface Wnt display and signaling of wild-type HEK293F and HEK293 expressing hFzd5_CRD_-1TM (top). Wnt expression detected by protein C (PrC) epitope tag on Wnt (bottom). (B) Ribbon models of XWnt8-mFzd8_CRD_ (PDB 4F0A) and hWnt3-mFzd8_CRD_ (PDB 6AHY) (top). The NC-linker is missing in the XWnt8 crystal structure, likely reflecting its flexibility. Cell surface expression and LRP6_E1E2_ tetramer binding on hFzd5_CRD_-1TM cell line in the absence or presence of wild-type XWnt8 or XWnt8_ΔNC_ (bottom). (C) Improvements in LRP6_E1E2_ tetramer binding to the XWnt8 libraries during selection compared to the wild-type XWnt8 single clone. Data were analyzed with FCS Express 7. (D) LRP6_E1E2_ monomer binding to the haXWnt8 and wild-type XWnt8 (left) and hahWnt8 and wild-type hWnt8 (right). The squares and dots indicate the means of experimental values, and the error bars represent standard deviation (SD). Data were analyzed with Prism 9. The NC-linker sequences are presented on top of the binding curves. (E) Top-flash signaling assays with recombinant wild-type XWnt8, haXWnt8, and next-generation surrogate (NGS) Wnt (Miao et al., 2020) (left panel) in the absence or presence of R-spondin 2. With two-fold serial dilution, the agonist concentrations were 0.39 nM to 50 nM for XWnt8s and 0.016 nM to 2 nM for NGS Wnt, respectively. The bar graphs show the means of experimental values, and the error bars represent SD. Data were analyzed with Prism 9. Dose-response experiments were performed in triplicate, and representative data were shown from two independent experiments.

After three rounds of fluorescence-activated cell sorting (FACS) selection for hLRP6_E1E2_ binding (Figure 1A and Figure 1—figure supplement 1B), we isolated five XWnt8 clones with improved hLRP6_E1E2_ tetramer binding relative to wild-type XWnt8 (Figure 1—figure supplement 2A). After screening for recombinant expression, we tested the XWnt8 variant with the highest expression (high-affinity XWnt8, haXWnt8) for hLRP6_E1E2_ monomer binding on cells, which had acquired six amino acid substitutions (Figure 1D). The haXWnt8 variant had at least 6-fold improved binding affinity (EC_50_) relative to wild-type XWnt8 (Figure 1D left panel) and grafting this linker into the corresponding region of human Wnt8 (hWnt8) yielded a similarly higher affinity (Figure 1D right panel and Figure 1—figure supplement 2B). To determine the functional outcomes of enhanced LRP6 binding, we performed super TOPflash Wnt reporter assays using recombinant XWnt8 or haXWnt8 in comparison with surrogate (NGS) Wnt (Miao et al., 2020) (Figure 1E). haXWnt8 showed more potent β-catenin signaling than the wild-type XWnt8 both in the absence and presence of R-spondin 2. In addition, we noticed that the surrogate Wnt showed a substantially better EC_50_ than the Wnts due to its high affinity but with a lower maximum effect (E_max_), which is likely due to a different, less optimal Fzd-LRP6 heterodimer receptor geometry than induced by Wnt.

### Reconstitution of the soluble Wnt ternary complex and structural analysis

To prepare the soluble ternary complex, we co-cultured S2 cells expressing haWnt8-mFzd8_CRD_ and hLRP6_E1E2,_ and the complex was purified by a series of affinity and size-exclusion chromatography steps. The peak fractions corresponding to the 1:1:1 haXWnt8-mFzd8_CRD_-hLRP6_E1E2_ complex were subjected to cryo-EM analysis (Figure 2A-D, Table 1, and Figure 2— figure supplement 1). 2D class averages of the complex revealed the central Wnt bridging the small globular mFzd8_CRD_ and the tandem β-propellers of hLRP6_E1E2_ (Figure 2B). The 3D reconstruction yielded the 3.8 Å nominal resolution map, revealing the stereotypical Wnt-Fzd_CRD_ binding mode as previously reported (Hirai et al., 2019; Janda et al., 2012) (Figure 2C). hLRP6_E1E2_ appears to “cap” XWnt8 at the top of the structure by engaging two sites on XWnt8 distal from the Fzd binding site. Site A is formed by the XWnt8 N-terminal loop, and Site B is formed by the NC-linker, which insert into the E1 and E2 funnels in the center of the hLRP6_E1E2_ β-propellers, respectively (Figure 2D). Both binding motifs in XWnt8 were not resolved in the previously reported XWnt8-mFzd8_CRD_ crystal structure (Janda et al., 2012), suggesting they are disordered in the absence of LRP6.

**Table 1.** Cryo-EM data collection and modeling.

| haXWnt8-mFzd8<sub>CRD</sub>-hLRP6<sub>E1E2</sub><br>(EMDB EMD-26989, PDB 8CTG) |  |
| --- | --- |
| **Data collection and processing** |  |
| Nominal magnification | 29,000x |
| Calibrated magnification | 58,680x |
| Voltage (kV) | 300 |
| Electron exposure (e/Å<sup>2</sup>) | 55 |
| Defocus range (μm) | -1.0 to -2.0 |
| Pixel size (Å) | 0.8521 |
| Symmetry imposed | C2 |
| Initial “particle” images (no.) | 3,840,364 |
| Final particle images (no.) | 82,235 |
| Map resolution (Å) | 3.8 (4.5) |
| FSC threshold | 0.143 (0.5) |
| Map resolution range (Å) | 4.4 to 7.2 |
| FSC threshold | 0.5 |
| **Refinement** |  |
| Initial models used (PDB codes) | 4F0A and 3S94 |
| Model resolution (Å) | 9.7 (4.4) |
| FSC threshold | 0.5 (0.143) |
| Map sharpening method | deepEMhanceer (visualization)<br>uniform B-factor of 200 Å<sup>2</sup> (blurred, refinement) |
| **Model composition** |  |
| Non-hydrogen atoms | 5,100 |
| Protein residues | 1,006 |
| Ligands | PAM |
| *B* factors (Å<sup>2</sup>) |  |
| Protein | 267.4 |
| Ligand | 551.1 (PAM) |
| **R.m.s. deviations** |  |
| Bond lengths (Å) | 0.004 |
| Bond angles (°) | 0.975 |
| **Validation** |  |
| MolProbity score | 2.83 |
| Clashscore | 12.46 |
| Poor rotamers (%) | 6.90 |
| **Ramachandran plot** |  |
| Favored (%) | 88.53 |
| Allowed (%) | 9.66 |
| Disallowed (%) | 1.81 |

**Figure 2.**
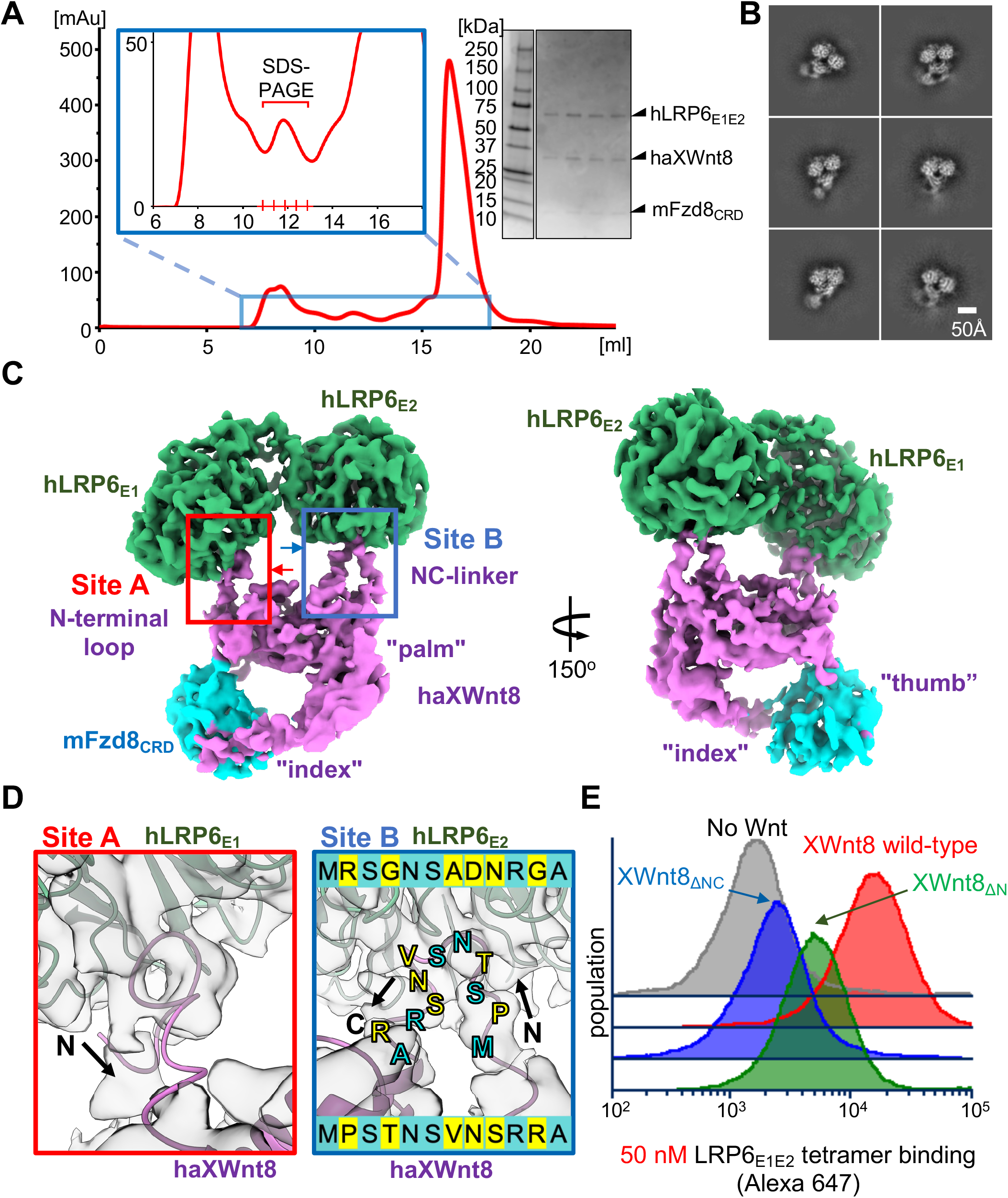
Reconstitution and cryo-EM analysis of haXWnt8-mFzd8_CRD_-hLRP6_E1E2_. (A) Size-exclusion chromatography profile and SDS-PAGE of haXWnt8-mFzd8_CRD_-hLRP6_E1E2._ (B) Representative 2D class averages of haXWnt8-mFzd8_CRD_-hLRP6_E1E2._ (C) 3D reconstruction of haXWnt8-mFzd8_CRD_-hLRP6_E1E2_ colored in purple (haXWnt8), cyan (mFzd8_CRD_), and green (hLRP6_E1E2_). (D) Closed-up of the Site A and Site B interfaces viewed from the red and blue arrows indicated in the panel c left. The cryo-EM map (transparent gray) is superimposed on the ribbon model of the complex (LRP6, green; haXWnt8, purple). The NC-linker shows continuous cryo-EM density, while N-terminal region does not. Map contour levels are set to 0.32 on ChimeraX. The sequence of the engineered haXWnt8 NC-linker is displayed at the bottom of the right panel and on the structure, with the wild-type sequence shown at the bottom. (E) LRP6_E1E2_ tetramer binding to hFzd5_CRD_-1TM cells with or without expression of wild-type XWnt8, XWnt8_ΔN_, or XWnt8_ΔNC_, showing that both the N-terminal loop and NC-linker contribute to the LRP6 binding. The tetramer concentration was 50 nM and four-fold lower than the experiment shown in Figure 1B. Figure 2—source data 1 | The uncropped SDS-PAGE of the size-exclusion chromatography fractions of the haXWnt8-mFzd8CRD-hLRP6_E1E2_ complex. Boxes indicate the region shown in Figure 2A.

The presumed flexibility of the unusual interaction mode seen, as compared to a more typical rigid protein-protein interface that buries large amounts of surface area, has limited the quality of our cryo-EM map, precluding placement of amino acid side chains for the XWnt8 N-terminus and NC-linker. While the N-terminal interaction is particularly tenuous, the NC-linker mainchain clearly shows that the central region of the loop inserts into the funnel. It seems likely that even when in complex, hinge flapping at the XWnt8-hLRP6_E1E2_ junction is an inherent feature of this interaction.

The observed Site A interaction mediated by the XWnt8 N terminus was surprising, so we asked if it was merely an adventitious, but energetically inert, interaction by virtue of its proximity to the NC-linker interaction with E2, or if it contributed to the overall binding free energy in forming the complex. There is precedent for Wnt N-termini being important for function in that Tiki family metalloproteases are Wnt inactivating enzymes that cleave N-terminal loops of Wnts (Zhang et al., 2016, 2012). As a matter of fact, Tiki inactivates both Wnt3a and XWnt8. Using our mammalian Wnt display system, we find that deleting the hydrophobic N-terminal loop (XWnt8_ΔN_) reduced LRP6 binding similar to XWnt8_ΔNC_ (Figure 2E). Since the effect was less pronounced than deleting the NC-linker, the NC-linker clearly plays the major role in driving the Wnt-LRP6 association, with the N-terminal loop being supportive, consistent with the less defined, discontinuous N-terminal cryo-EM density observed. Unlike most Wnts, Wnt8s have shorter N-terminal sequences preceding the N-domain, lacking a subdomain containing a disulfide bond (Hirai et al., 2019; Nygaard et al., 2021; Zhong et al., 2021). However, with a few exceptions, Wnts have comparably long disordered loops with hydrophobic residues, reinforcing the shared Wnt-LRP6 binding mechanism in canonical β-catenin signaling.

### Generality of NC-linker to mammalian Wnts

We wished to determine the generality and modularity of the NC-linker of XWnt8 and mammalian Wnts. The NC-linker is one of the least conserved amino acid stretches across different Wnt proteins, but it is highly conserved across species for a particular Wnt (Figure 3—figure supplement 1). Using human Wnt1 as a model system, we performed alanine scanning mutagenesis on the NC-linker analogous to that of XWnt8 to assess its functional role in mediating Wnt/β-catenin signaling. We transfected HEK293 TOPbrite luciferase reporter cells (Nile et al., 2018) with plasmids encoding Wnt1 mutants with pairs of alanine mutations. Despite comparable expression, most Wnt1 variants were less active in inducing β-catenin signaling compared to wild-type Wnt1, with mutants 65, 70, 73, and 76 having the most impaired signaling activities (Figure 3A and Figure 3—figure supplement 2). Thus, the NC-linker of human Wnt1 also appears to be a critical mediator of Wnt signaling activity, and this likely extends to other Wnt family members and species.

**Figure 3.**
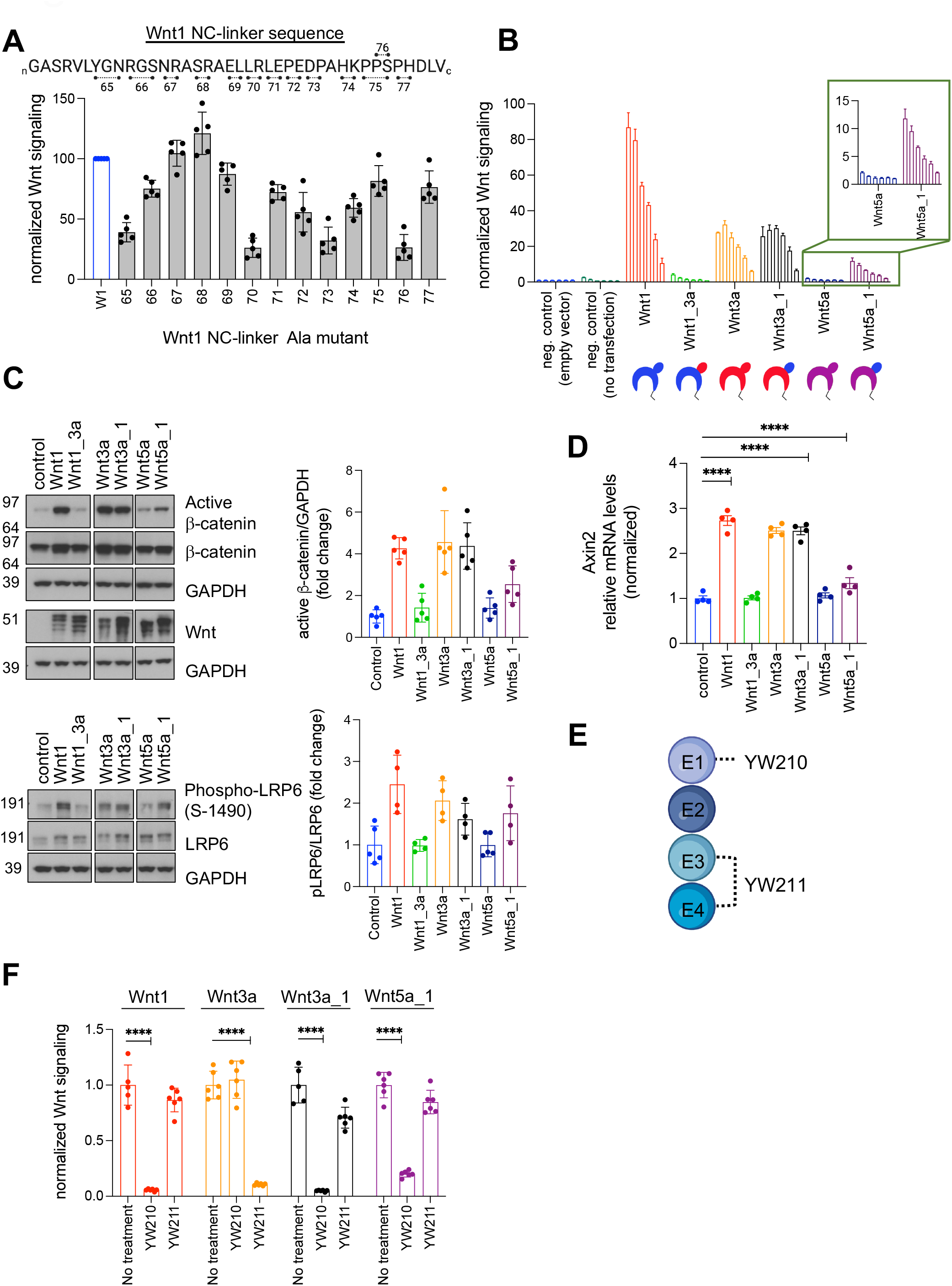
Exchange of the NC-linker between Wnt proteins modifies the signaling characteristics of their parental Wnt. (A) Representative TOPbrite dual-luciferase reporter assay showing normalized Wnt signaling induced by various Wnt1 NC-linker alanine mutants from five biological replicates. Sequences are provided in Figure S5. Bar and error bar represent the mean and SD of five technical replicates. (B) Representative TOPbrite dual-luciferase reporter assay measuring Wnt signaling with overexpression of Wnts and Wnt chimeras. Signaling was normalized to the negative control (empty vector). Bar and error bar represent the mean and SD of three technical replicates from one of three biological replicates. (C) Representative Western blot showing β-catenin and LRP6 levels with overexpression of Wnts and Wnt chimeras. Quantification is provided next to the blots. Bar and error bar represent the mean and SD of four to five biological replicates. (D) quantitative PCR analysis of Wnt target gene *Axin2* with overexpression of Wnts and Wnt chimeras. The expression was normalized to the negative control in each group. Bars represent the means and standard error of the mean of four biological replicates. (E) Schematic representation of LRP6 domains and domain-specific bound antibodies, YW210 and YW211. (F) Representative TOPbrite dual-luciferase reporter assay showing the signaling characteristics of Wnt1, Wnt3a and the Wnt1-mimicking chimeras, Wnt3a_1 and Wnt5a_1. The signaling was normalized to the negative control (without any treatment) in each group. Bar and error bar represent the means and SD of six technical replicates. (****p<0.0001, one-way ANOVA with Tukey’s multiple comparison test) from one of three biological replicates. Figure 3—source data 1 | The raw western blot data showing β-catenin and LRP6 levels with overexpression of Wnts and Wnt chimeras. The boxes indicate the region shown in Figure 3C.

### The NC-linker controls Wnt-LRP6 domain specificity

To investigate the modularity of the NC-linker in mediating LRP6 interactions by different Wnts, we designed three chimeric Wnt constructs in which the NC-linker was swapped between different human Wnt proteins (Figure 3B). We chose Wnt1, Wnt3a, and Wnt5a as templates for NC-linker swapping because they represent canonical (Wnt1 and Wnt3a) and non-canonical (Wnt5a) Wnts and interact with LRP6_E1E2_ (Wnt1) and LRP6_E3E4_ (Wnt3a) (Bourhis et al., 2010; Gong et al., 2010). Of note, since no structures of Wnt1 and Wnt5a were available, the designs of the chimeras, particularly the NC-linker boundaries, were not informed by structure and are therefore approximations. We expect that these structural imprecisions of the NC-linker grafts in the chimeric molecules will quantitatively influence their activities, and also that some loops will resist grafting due to structural differences between the NC-linker attachment sites to the Wnt globular core. With these caveats in mind we undertook a range of grafts and assessed their activities. As expected, the canonical Wnt proteins Wnt1 and Wnt3a induced β-catenin signaling in TOPbrite assays, whereas the non-canonical Wnt5a did not induce reporter expression above background (Grumolato et al., 2010) (Figure 3B). Grafting the NC-linker from the E1E2-binding Wnt1 into the E3E4-binding Wnt3a (termed Wnt3a_1) did not markedly alter the activity of wild-type Wnt3a, suggesting that we successfully transferred LRP6_E1E2_ E1E2 specificity from Wnt 1 to Wnt3a. However, the reverse engineered Wnt1_3a chimera was inactive. This could reflect the fact that the NC-linker was disordered in the XWnt8-mFzd8_CRD_ crystal structure, whereas the Wnt3a NC-linker was ordered into a β-hairpin. As a result, grafting the Wnt3a NC-linker may have less tolerance for different structural contexts. Consistent with this, grafting the Wnt1 NC-linker into Wnt5a rendered the Wnt5a_1 chimera able to induce β-catenin signaling (Figure 3B), functionally turning a classical non-canonical Wnt into a canonical Wnt. Consistently, Wnt1, Wnt3a, Wnt3a_1, and Wnt5a_1 but not Wnt5a and Wnt1_3a induced LRP6 phosphorylation, the accumulation of active (non-phosphorylated) β-catenin (Figure 3C) and Axin2 transcription (Figure 3D), all characteristic hallmarks of canonical Wnt signaling. Grafting the Wnt3a NC-linker into Wnt5a (Wnt5a_3a) did not result in activity.

To further interrogate the LRP6_E1E2_ *versus* LRP6_E3E4_ binding specificity of the Wnt chimeras, and the role of the NC-linker in determining binding preferences, we leveraged two LRP6 domain-specific antibody fragments (Fab): Fab YW210.09 (named YW210 hereafter) binds predominantly to LRP6 E1 domain, and Fab YW211.31.62 (named YW211 hereafter) binds to LRP6 E3E4 domain (Bourhis et al., 2011; Gong et al., 2010) (Figure 3E). As expected, YW210 inhibited Wnt1-induced signaling and YW211 inhibited Wnt3a-induced signaling in luciferase assays, consistent with the reported LRP6 domain specificity (Figure 3F and Figure 3—figure supplement 3). However, the activities of the Wnt3a_1 and Wnt5a_1 chimeras were inhibited by YW210, and not by YW211, demonstrating that the Wnt1 NC-linker directs the binding of these chimeras to LRP6_E1E2,_ mimicking Wnt1 signaling characteristics. In summary, these results suggest a critical role of the Wnt NC-linkers in determining canonical and non-canonical Wnt signaling activity and LRP6 domain specificity, and that the signaling characteristics of Wnts can be re-engineered by transferring the NC-linker. While not all grafts were successful, we think the modularity of the NC-linker demonstrated for Wnt1 is likely representative for most Wnts.

### Synthetic peptides mimicking NC-linker inhibit Wnt activity

To further support the role of the NC-linker as a critical determinant of LRP6 domain specificity, we investigated the binding specificities of the isolated NC-linkers and their ability to interfere with Wnt activity. Superposition of the three available wild-type Wnt structures (Chu et al., 2013; Hirai et al., 2019; Janda et al., 2012) revealed two loosely structurally conserved ‘interface motifs (Bazan et al., 2012; Hirai et al., 2019)’ that flank the variable NC-linkers (Figure 4—figure supplement 1A,B). We designed cyclic peptides mimicking the NC-linkers (termed W.cys) of four different Wnts and engineered cysteine residues at two conserved positions within the interface motifs that are within optimal distance (Cβ - Cβ distance of ∼4.0 Å) for disulfide bond formation (Figure 4A and Figure 4—figure supplement 1B). In an ELISA experiment, we observed strong binding of W1.cys, W2b.cys and W7a.cys to LRP6_E1E2_ (Figure 4B left panel), and of W3a.cys to LRP6_E3E4_ (Figure 4B right panel and Table 2). This is consistent with previously reported Wnt-LRP6 domain specificity^14^. We further demonstrated that W7a.cys, but none of the other NC-linker peptides, bound Reck (Figure 4—figure supplement 2A), consistent with an earlier report (Cho et al., 2019), proposing a more general function of the NC-linker in recruiting co-receptors. The weaker interaction between W3a.cys and LRP6_E1E2_ can be attributed to “sticky” non-specific binding as it could not be outcompeted by anti-LRP6_E1_ YW210 (Figure 4C). Despite the inherent flexibility of isolated peptides, W1.cys, W2b.cys and W7a.cys, but not W3a.cys, inhibited Wnt1-mediated β-catenin signaling in luciferase assays without any observed toxicity on cells (Figure 4D and Figure 4—figure supplement 2B), presumably by competing with Wnt1 for LRP6_E1E2_ binding. The lack of activity of the Wnt3a peptide is consistent with its binding ability to LRP6_E3E4_. Taken together, the overall consistency in LRP6 domain specificity of isolated NC-linker peptides and full-length Wnt proteins, as well as the ability of the peptides to interfere with Wnt activity, further supports the critical role of the NC-linker in determining LRP6 domain specificity.

**Table 2.** EC_50_ and R_max_ values determined by ELISA.

| Peptide | EC<sub>50</sub> (μM),<br>E1E2 | R<sub>max</sub> (a.u.), E1E2 | EC<sub>50</sub> (μM), E3E4 | R<sub>max</sub> (a.u.), E3E4 |
| --- | --- | --- | --- | --- |
| W1.cys | 0.067 ± 0.047 | 2.22 ± 0.31 | - |  |
| W2b.cys | 0.018 ± 0.015 | 2.06 ± 0.24 | - |  |
| W3a.cys | 0.22 ± 0.092 | 0.99 ± 0.53 | 0.19 ± 0.16 | 1.47± 0.29 |
| W7a.cys | 0.04 ± 0.018 | 2.13 ± 0.09 | - |  |

**Figure 4.**
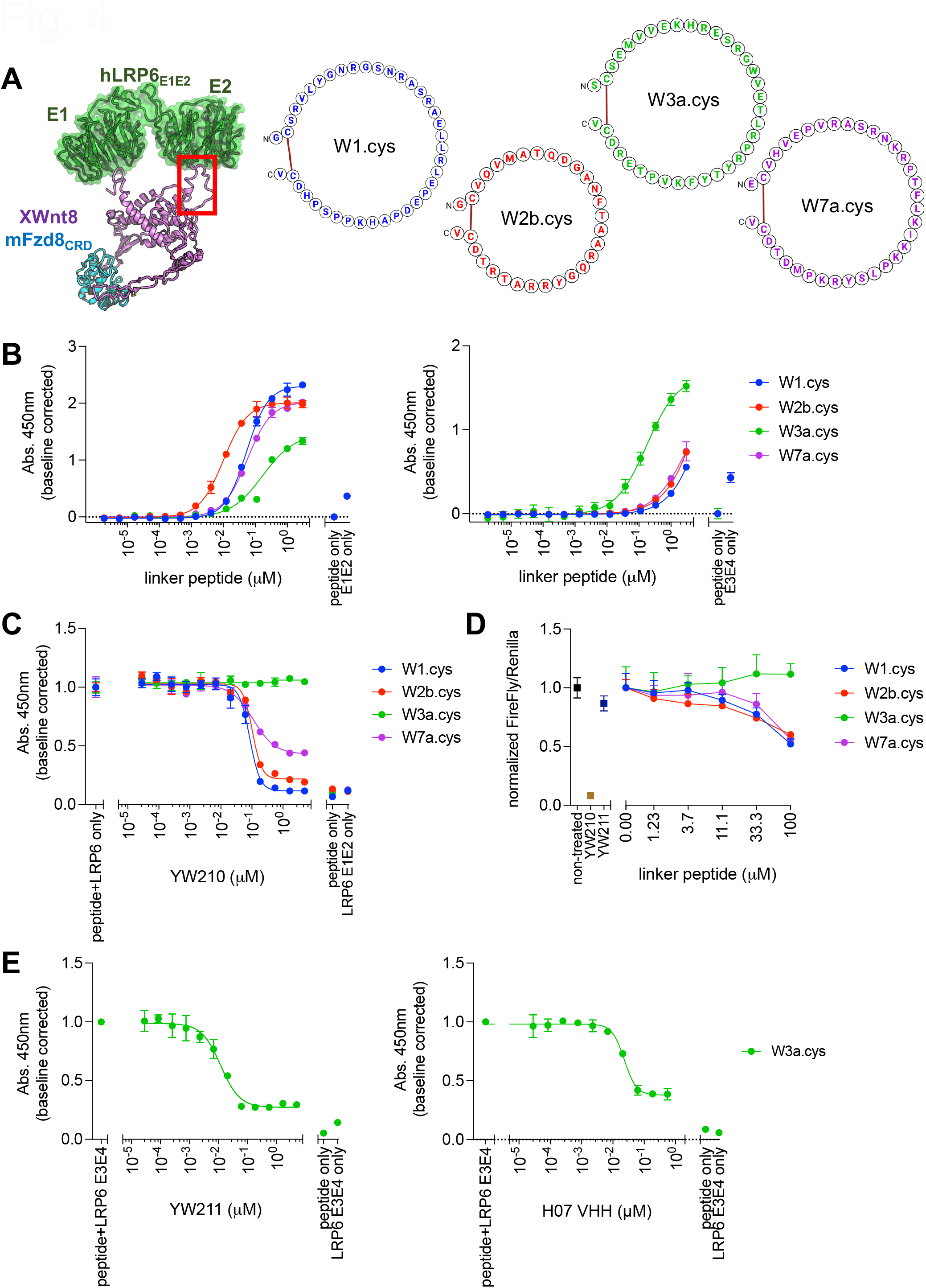
Synthetic Wnt linker peptides show LRP6 domain-specific binding. (A) A ribbon-and-surface model of haXWnt8-mFzd8_CRD_-hLRP6_E1E2_ colored in purple (haXWnt8), cyan (mFzd8_CRD_), and green (hLRP6_E1E2_) (left). The sequences of W1.cys, W2b.cys, W3a.cys, W7a.cys and W10b.cys are portrayed in the schematic form showing the constrained peptides by a disulfide bond (left). (B) Representative ELISA binding assay of Wnt.cys peptides to LRP6_E1E2_ (left) and LRP6_E3E4_ (right). The signals from the binding of various Wnt.cys peptides to the assay plate were subtracted from their binding signals to LRP6. Data are shown as the means with SD of three technical replicates from one of three biological replicates. (C) Dose-dependent inhibition of Wnt1-mediated signaling by W1.cys, W2b.cys and W7a.cys. Data are shown as the means with SD of three technical replicates from one of three biological replicates. (D) Representative ELISA competition assays of LRP6 E1E2-binding Wnt.cys with Fab YW210 and (E) LRP6 E3E4-binding Wnt.cys with Fab YW211 and H07 VHH. Only the peptides showing the binding to each functional domain of LRP6 were tested in the competition ELISA. Data are shown as the means with SD of three technical replicates from one of three biological replicates.

To further map the binding region of the NC-linker peptides on LRP6, we performed a competition ELISA with YW210, YW211, and H07, a single-domain antibody fragment (VHH) that binds to the LRP6 E3 domain (Fenderico et al., 2019). YW210 competed strongly with W1.cys and W2b.cys, and partially with W7a.cys for LRP6_E1E2_ binding (Figure 4C). Given the relatively large size of Fabs, it likely sterically masked neighboring binding epitopes on LRP6_E2_. Furthermore, YW211 and H07 VHH both competed with W3a.cys for binding to LRP6_E3_ (Figure 4E), suggesting that the Wnt3a NC-linker binds to the E3 domain of LRP6.

hLRP6_E1E2_ and hLRP6_E3E4_ are relatively conserved regarding their funnel-like cavity at the center of the β-propellers, hydrophobicity and electrostatic property of β-propellers, and the ∼40 Å distance between the proximal funnels (Figure 5—figure supplement 1), consistent with a shared two-site Wnt binding mechanism by both LRP6_E1E2_ and LRP6_E3E4_. Our H07 VHH blocking data shows that E3 domain blockade prevents Wnt3a binding, localizing the NC-linker to the LRP6 E3 domain, while the NC-linker of XWnt8 and Wnt1 binds to the LRP6 E2 domain. This structure-function data now allows us to orient how E1E2-binding versus E3E4-binding Wnts engage LRP6 through their NC-linkers and N termini. We propose that, like XWnt8, E1E2-binding Wnts use their N termini to bind E1 and NC-linker to bind E2, whereas E3E4-binding Wnts, represented by Wnt3, use their N termini to bind E4 and their NC-linker to bind to E3. Thus E1E2-binding versus E3E4-binding Wnts appear to bind to LRP6 domains in ‘reverse’ orientations to accommodate the NC-linker specificity (Figure 5A). Whether all Wnts use their N termini to bind to the neighboring propeller funnel and orient themselves is not addressed by our data, but the perfect match of 40 Å distance between the N-termini and NC-linker on Wnts with the 40 Å distance between funnels on neighboring propellers suggests this is likely.

**Figure 5.**
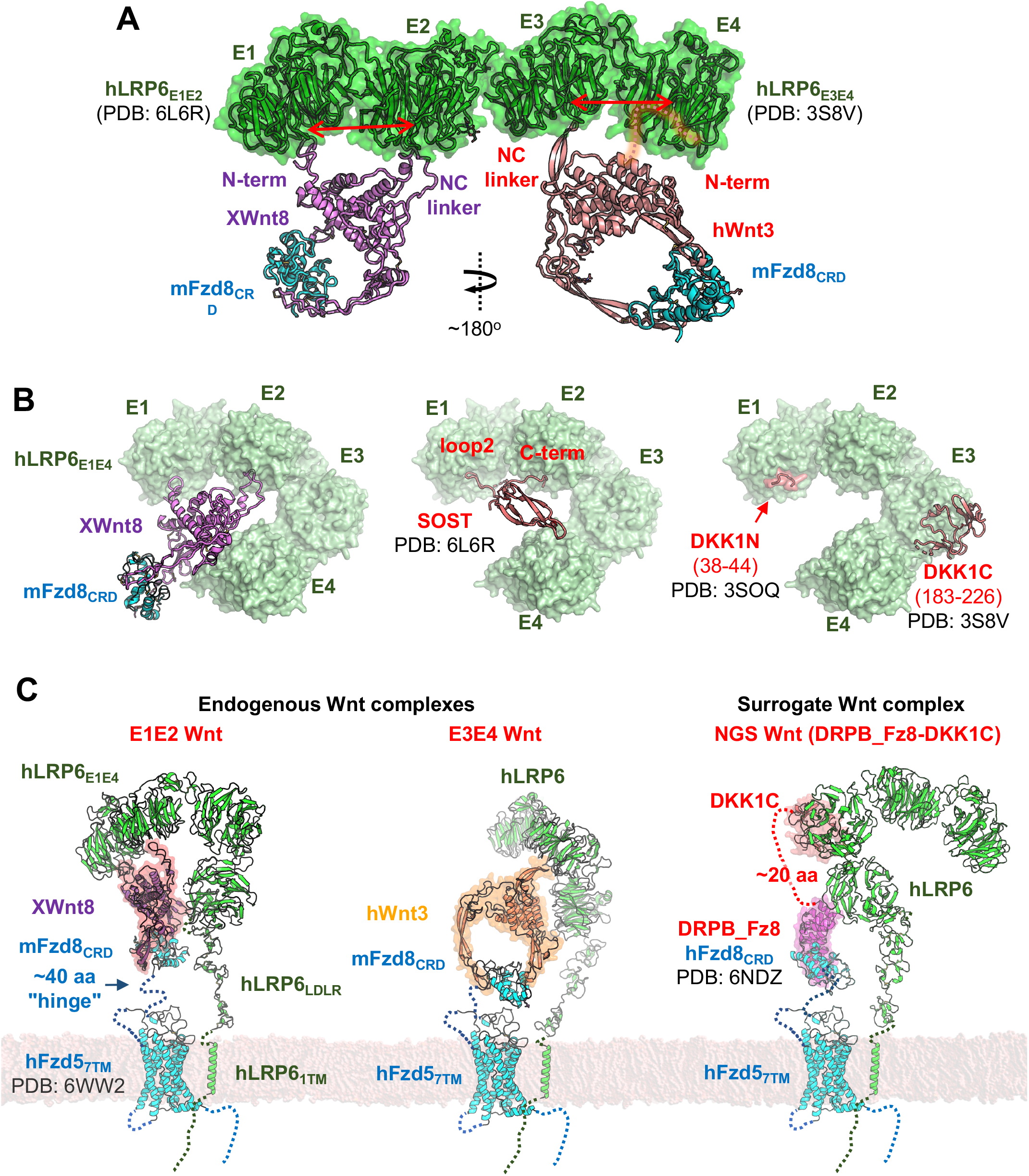
Modeling initiation and inhibition of Wnt signaling. (A) Binding modes of E1E2-binding Wnt8 and E3E4-binding Wnt3 to LRP6. Crystal structures of LRP6_E1E2_ and LRP6_E3E4_ are placed linearly, and mFzd8_CRD_-XWnt8 and mFzd8_CRD_-hWnt3 are anchored to the propeller funnels in the orientation suggested by the competition assays. LRP6, mFzd8, XWnt8, and hWnt3 are colored in green, cyan, purple, and pink, respectively. (B) The structural models of hLRP6_E1E4_ binding to haXWnt8-mFzd8_CRD_ (left), sclerostin (center), or DKK1 (right). (C) Schematic for the transmembrane LRP6-Fzd5 signaling complex with endogenous Wnts or Surrogate Wnt (Miao et al., 2020).

## Discussion

The structure of the ternary haXWnt8-mFzd8_CRD_-hLRP6_E1E2_ complex resolves the basis for co-receptor binding by Wnts, revealing a modular determinant that not only controls LRP5/6 domain specificity, but also appears to differentiate the non-canonical Wnt5a and explain its co-receptor independence. The signaling complex structure further elucidates the mechanism of LRP5 gain-of-functions mutations (including D111Y, R154M, G171V/R, N198S, A214V/T and T253I - which correspond to D98, R141, G158, N185, A201, and T240 in LRP6) that are associated with high bone mass (HBM) disease and the significant increase in bone strength and thickness in affected patients (Babij et al., 2003; Gregson and Duncan, 2020). These HBM mutations are located on the top surface of the E1 β-propeller of LRP5, and overlap with the binding epitopes of Wnt antagonists sclerostin (SOST) (Winkler et al., 2003) or Dickkopf (DKK) (Glinka et al., 1998; Krupnik et al., 1999) (Figure 5B). Consistent with the notion that the Wnt NC-linker-LRP6 E2 interaction is the main driver for binding, HBM mutations impaired the binding of SOST and DKK1, but with a minimal effect on Wnt9b in functional assays, resulting in the selective loss in affinity for Wnt signaling antagonists (Ai et al., 2005; Bourhis et al., 2011; Semenov and He, 2006).

The ECD of LRP5/6 consists of four YWTD β-propellers flanked by EGF-like domains, E1-E4 (Figure 5B left panel), followed by three low-density lipoprotein receptor type A domains, and exists in a range of bend conformations (Matoba et al., 2017). The ECD interacts with Wnt proteins and antagonists, including SOST and DKK, to regulate signaling. While SOST antagonizes LRP6_E1E2_-binding Wnts including Wnt1, Wnt2, and Wnt9b, but not LRP6_E3E4_-binding Wnts including Wnt3a (Kim et al., 2020), DKK1 inhibits both the Wnt groups (Bourhis et al., 2010; Sima et al., 2016). Our cryo-EM structure of the ternary haXWnt8-mFzd8_CRD_-hLRP6_E1E2_ complex now provides insights into the precise mechanisms of differential Wnt inhibition by SOST and DKK.

SOST is a secreted glycoprotein predominantly expressed by osteocytes and inhibits bone formation and regeneration by antagonizing Wnt/β-catenin signaling (Winkler et al., 2003). SOST engages the top surfaces of the E1 and E2 β-propellers with its loop 2 and the C-terminal tail (Bourhis et al., 2011; Kim et al., 2020), respectively, reminiscent of the tandem binding mode of haXWnt8. SOST occludes both Site A and B of the haXWnt8-LRP6 interfaces (Figure 5B middle panel), demonstrating that SOST antagonizes Wnt signaling predominantly through the direct competition with Wnts for LRP6 binding. This competitive tandem binding mode of SOST and Wnt is further supported by functional studies showing that both loop 2 and the C-terminal tail of SOST are critical for inhibiting β-catenin signaling mediated by Wnt1, Wnt2, and Wnt9b (Kim et al., 2020).

DKK comprises another class of four broadly expressed Wnt antagonists (Glinka et al., 1998; Krupnik et al., 1999). The N- and C-terminal fragments of DKK1, DKK1N (Bourhis et al., 2011) and DKK1C (Cheng et al., 2011), bind to the funnel surface of the LRP6 E1 and E3 β-propellers, respectively (Figure 5B right panel). The binding site of DKK1N overlaps with that of the Wnt N-terminal loop on E1, but not with the binding site of the NC-linker on E2, which is the major driver of the Wnt-LRP6 interaction. Upon binding of DKK1 to LRP6 through this unusual distant two-site binding, the LRP6 ECD adopts a compact conformation, bringing the E1E2 and E3E4 domains into close-proximity in a face-to-face orientation (Matoba et al., 2017). Together with our structure, this demonstrates that DKK1 inhibits Wnt binding to LRP6_E1E2_ and LRP6_E3E4_ by directly competing for LRP6 binding as well as through steric occlusion of Wnts from the compact ECD fold. Understanding the molecular mechanisms by which endogenous Wnt binding proteins and disease mutations alter Wnt signaling facilitate the development of more precisely designed therapeutic modalities to potentiate or inhibit Wnt signaling for regenerative medicine or cancer therapy, respectively.

Wnt proteins require the evolutionary conserved eight-pass transmembrane protein WLS for their secretion into the extracellular space. Wnts bind to WLS in the endoplasmic reticulum, and the complex is then shuttled through the Golgi to the cell membrane. Recent cryo-EM structures of Wnt3a and Wnt8 bound to WLS (Nygaard et al., 2021; Zhong et al., 2021) revealed that multiple Wnt loops and hairpins mediate WLS binding through an induced fit mechanism, enabled by the high flexibility of these Wnt regions (Figure 5—figure supplement 2A). This raises the possibility of a regulatory role of the Wnt intrinsic flexibility in mediating the interaction with different binding partners. For instance, the N-terminal loop of XWnt8/Wnt8 (and Wnt3/Wnt3a) that engages the top surface of the E1 β-propeller in our cryo-EM structure is flexible and unresolved in the crystal structures when bound to the Fzd_CRD_ but is structured when bound to the WLS cargo receptor and packs tightly against the Wnt globular domain (Hirai et al., 2019; Janda et al., 2012; Nygaard et al., 2021; Zhong et al., 2021) (Figure 5—figure supplement 2B). The N-terminus might dislodged from the Wnt core upon binding to Fzd_CRD_, which may result in being more accessible by LRP6. However, this allosteric “priming” of Wnt by Fzd_CRD_ binding is only supported by indirect structural evidence.

The ternary XWnt8-mFzd8_CRD_-hLRP6_E1E2_ complex structure, together with the structure of the Fzd transmembrane regions (Tsutsumi et al., 2020; Yang et al., 2018), and previous mechanistic insights into heterodimeric Wnt activation (Miao et al., 2020; Tsutsumi et al., 2020) allow us to propose a structural model of the transmembrane Wnt signaling complex for both LRP6 E1E2 and E3E4 binding Wnts (Figure 5C). Given the different topologies of these signaling complexes it is tempting to speculate that this could affect signaling strength. We have recently shown that bispecific Wnt “surrogates” composed of a Fzd_CRD_ binding domain fused to DKK1C can activate β-catenin signaling by simply heterodimerizing Fzd and LRP6. However, the surrogates with different binding domains and linkers show great variations in signaling strength, suggesting that the geometry (*i*.*e*., orientation and proximity) of the two receptor chains within the signaling complex is a critical determinant of signaling characteristics. Furthermore, the geometry of the signaling complex induced by the Wnt surrogates and natural Wnt ligands differs greatly, possibly yielding the differential E_max_. Thus, the haWnt8-mFzd8_CRD_-hLRP6_E1E2_ structure provides a blueprint to rationally improve the Wnt surrogates by modulating receptor complex geometry as has been done in homo- and heterodimeric cytokine receptors (Mohan et al., 2019; Yen et al., 2022), facilitating more effective Wnt surrogates for regenerative medicine.

## Materials and Methods

### Mammalian cell line for Wnt display

To display functional Wnts on cells, we engineered HEK293F cells (Thermo Fisher) to saturate the cell surface with hFzd5_CRD_ (residues 27-155) in a manner analogous to scFv displayed on mammalian cells (Chesnut et al., 1996). The N-terminal HA-tagged hFzd5_CRD_ was fused to the membrane-proximal region and transmembrane domain of gorilla Platelet-derived growth factor receptor-β (PDGFR-β, residues 513-561) with a linker containing Myc tag (PG**SGEQKLISEEDL**GNGNGNGNGNGNGNGNGNGNGNG), and cloned into the lentiviral gene ontology (LeGO) vector which has EF-1α promoter and Igκ signal peptide (Weber et al., 2010), termed pTether hFzd5_CRD_-1TM. For lentivirus packaging, Lenti-X 293T cells (Takara Bio) were transfected with pTether hFzd5_CRD_-1TM, pMD2.G (Addgene 12259), and psPAX2 (Addgene 12260) vectors using FuGENE HD (Promega), and the cells were cultured at 37°C for 3 days. Lenti-X 293T cells were maintained in Dulbecco’s Modified Eagle Medium (DMEM, Gibco) supplemented with 10% (v/v) Fetal Bovine Serum (FBS, Sigma) and GlutaMAX (Gibco), or “complete DMEM”, at 37°C and 5% CO_2_. The media was exchanged with FreeStyle 293 media (Thermo Fisher) prior to the transfection, and the supernatant containing the lentivirus was harvested and used immediately to infect HEK293F cells maintained in FreeStyle 293 media at 37°C and 5% CO_2_ with gentle agitation. After 3 days, cells were stained using Alexa Fluor 647 Conjugated HA-Tag (6E2) Mouse mAb (Cell Signaling 3444) and analyzed by an Accuri C6 Plus flow cytometer (BD Bioscience) to check the surface hFzd5_CRD_ expression, and high-expression clones were bulk sorted by FACS using SH800S Cell Sorter (Sony Biotechnology).

### Wnt display

To test Wnt display on the cell surface, *Xenopus laevis* Wnt8 (XWnt8) was expressed in the parental HEK293F cells and the hFzd5_CRD_-1TM cells before FACS sorting. Full-length XWnt8 was cloned into pcDNA3.1+ (Thermo Fisher) vector with endogenous signal peptide and C-terminal Protein C epitope. The cells were transiently transfected with the pcDNA3.1+ XWnt8 vector and stained by Alexa Fluor 647 conjugated Protein C antibody (PrC647; the antibody produced in-house from HPC-4 hybridoma, ATCC HB-9892, and labeled with Alexa Fluor 647 NHS Ester, Thermo Fisher), showing superior Wnt display on the hFzd5_CRD_-1TM cells. The flow cytometry data were analyzed using FCS Express 7 (De Novo Software).

To confirm that the lipidated Wnt stays on the cell which produced the molecule, the sorted hFzd5_CRD_-1TM cell line was transfected either with the XWnt8 or green fluorescent protein (GFP)-coding mammalian expression vector, and 24 hrs after transfection, the Wnt and GFP induced cells were mixed at 1:1 ratio and further shaken at 37°C for 24 hrs. Cell-surface Wnt was labeled with PrC647, and Wnt/GFP expression was checked by the flow cytometry. Wnt transfer to the GFP positive cells was estimated to be ∼5%.

To prepare stable cell lines expressing XWnt8 or hWnt8 with or without the deletion of the NC-linkers (residues 222-234 for XWnt8, and 221-233 for hWnt8, that were replaced by GSGS), and N-terminal loops (residues 23-32 for XWnt8), the constructs were cloned into the pLV-EF1a-IRES-Puro vector (Addgene 85132) with N-terminal HA signal peptide and C-terminal FLAG tag. The lentiviruses were prepared and added to the hFzd5_CRD_-1TM cell line, and 2 days after infection, the transduced cells were selected by adding 1 μg/ml puromycin (Gibco) and incubated for 2 more days. Wnt display was confirmed by mouse Monoclonal ANTI-FLAG M2-FITC antibody (FLAG-FITC, Sigma F4049), showing comparable display between the full-length and NC-linker truncated versions.

### Preparation of recombinant hLRP6_E1E2_ for cell staining

Like the majority of Wnts, XWnt8 and hWnt8 bind to the E1E2 module of human LRP6 (hLRP6_E1E2_). To recombinantly express hLRP6_E1E2_ and check Wnt8-hLRP6_E1E2_ interaction, hLRP6_E1E2_ (residue 20-629) was cloned into pAcGP67A (BD Biosciences) vector with C-terminal Biotin Accepter Peptide (BAP) and hexahistidine tags. *Spodoptera frugiperda* (Sf9) ovarian cells (ATCC CTL-1711) were maintained in Sf-900 III medium (Gibco) with 10% (v/v) FBS and GlutaMAX at 27°C with ambient CO_2_ and gentle agitation. P0 virus was produced by transfection with FuGENE HD and BestBac 1.0 Linearized Baculovirus DNA (Expression Systems) followed by a viral amplification to P1. The protein was expressed in *Trichoplusia ni* ovarian cells (Expression Systems) for 60-72 hrs maintained in ESF 921 Insect Cell Culture Medium (Expression Systems) at 27°C with ambient CO_2_ and gentle agitation. The protein was purified from culture supernatant with Ni-NTA (Qiagen), biotinylated with in-house purified BirA enzyme, purified over a Superdex 200 10/300 GL size-exclusion chromatography (SEC) column (Cytiva) equilibrated with HBS (10 mM HEPES-Na and 150 mM NaCl) and 2 mM CaCl_2_. The peak fraction was concentrated to ∼30 μM, aliquoted, and flash frozen. A frozen aliquot was thawed and mixed with Streptavidin (Thermo Fisher) for SDS-PAGE band-shift assay to confirm ∼100% biotinylation.

### hLRP6_E1E2_ tetramer staining of the Wnt-expressing cells

Biotinylated hLRP6_E1E2_ was tetramerized with Alexa 647 labeled streptavidin (SA647, streptavidin with a cysteine residue produced in house, and labeled with Alexa Fluor 647 C2 Maleimide, Thermo Fisher) and cells were stained with 30-200 nM hLRP6_E1E2_-SA647 tetramer (120-800 nM hLRP6_E1E2_ as monomer). The stained cells were washed with PFE (PBS, 2% (v/v) FBS, 2 mM EDTA) twice, and analyzed with the flow cytometer. The initial tests were performed with transient Wnt expression, and the tetramer staining was further confirmed for stable XWnt8 and hWnt8 cell lines described above. Clear tetramer staining was observed for the wild-type Wnt8s, but as suggested for hWnt3a (Chu et al., 2013; Hirai et al., 2019), the NC-linker deletion decreased hLRP6_E1E2_ binding to Wnt8s (Figure 1B and Figure 1—figure supplement 2), underscoring the premise for library creation on this linker. The flow cytometry data were analyzed using FCS Express 7.

### XWnt8 library construction

After confirming that the Wnt8s’ NC-linker mediates interaction with hLRP6_E1E2_, two XWnt8 libraries were constructed on the linker targeting 1) family conserved amino acid residues and 2) other amino acid residues (Figure 1—figure supplement 1A). In the parental pLV-EF1a-IRES-Puro XWnt8 plasmid, the XWnt8 gene was codon-optimized, and restriction enzyme sites were introduced at around E201 (EcoRI) and G247 (BamHI) with silent mutations (Figure 1—figure supplement 1A). The library genes were ordered from Eurofins Scientific as EXTREmers Long DNA and amplified by PCR for joining with the XWnt8 vector cut by the EcoRI and BamHI enzymes (New England Biolabs, NEB). Each library insert and the cut XWnt8 vector were mixed at 1:2 molar ratio, and the DNAs were assembled by NEBuilder HiFi DNA Assembly Master Mix (NEB) and purified with Zymo-Spin I Columns (Zymo Research) for electroporation to *E. coli*. The purified DNA was added to MegaX DH10B T1R Electrocomp Cells (Thermo) on ice, and the cells were placed in the ice-cold electroporation cuvettes (2 mm gap, Fisher Scientific) for transformation with MicroPulser Electroporator (BioRad). The cells were incubated with the provided Recovery Medium before plating. The efficiency was checked with serial dilution on Lysogeny broth (LB) plates supplemented with 100 μg/ml carbenicillin, and according to the titration result, the freshly electroporated and recovered *E. coli* libraries were plated on eight each of LB square bioassay dishes (Corning) supplemented with 100 μg/ml carbenicillin and incubated at 37°C overnight. Approximately 0.5 million transformants were obtained for each library. The *E. coli* colonies were collected with scrapers and the library plasmids were purified using Plasmid Maxi Kit (Qiagen). Lentiviral libraries were prepared as described above in complete DMEM and harvested using PEG-it Virus Precipitation Solution (System Biosciences). Lentiviral titers were checked using the hFzd5_CRD_-1TM cells at 2 ml scale based on Wnt expression, and 120 ml of 1 × 10^6^ cells/ml were used to have ∼1% cells infected. After 3 days of infection, the cells were stained by FLAG-FITC and the transduced cells were isolated by magnetic-activated cell sorting using Anti-FITC MicroBeads (Miltenyi) and LS columns (Miltenyi). The library cells were grown in FreeStyle 293 media, and once the total cell number reached >10 million for both library, 5 million cells each of FLAG-FITC+ library 1 and 2 were mixed to make the naïve XWnt8 library.

### XWnt8 library selection and the NC-linker chimera of hWnt8a

The naïve XWnt8 library was stained with 100 nM hLRP6_E1E2_-SA647 tetramer and 1:50 FLAG-FITC at 4°C for 1 hr, washed twice with PFE, passed through 70 μm Nylon Sterile Cell Strainers (Corning), and sorted using SH800S Cell Sorter. Top 7-8% clones based on XWnt8 expression (FLAG-FITC) and hLRP6_E1E2_ binding (SA647) were selected, then washed with and re-grown in FreeStyle 293 media. The following rounds of selections were performed in the same manner as the first round with increased stringency: the second selection was performed with 50 nM hLRP6_E1E2_ tetramer and the selection of top 7-8% clones, and the final, third round was done with 30 nM hLRP6_E1E2_ tetramer and ∼5% gating (Figure 1—figure supplement 1B). hLRP6_E1E2_ tetramer staining showed increased binding during the selection (Figure 1C). The genomes were then extracted from the post-round three library using QIAmp DNA Mini Kit (Qiagen), and the library regions were amplified by PCR and re-cloned into pLV-EF1a-IRES-Puro for sequencing, and hLRP6_E1E2_ binding experiments on the Wnt-expressing hFzd5_CRD_-1TM cells. All five single clones tested (named haXWnt8, haXWnt8a, haXWnt8b, haXWnt8c, haXWnt8d) showed improved binding to hLRP6_E1E2_ compared to the wild-type XWnt8 (Figure 1D and Figure 1—figure supplement 2).

The engineered NC-linkers from the XWnt8 library were grafted to hWnt8 to test if they improve the hLRP6_E1E2_ binding. The wild-type pLV-EF1a-IRES-Puro hWnt8 plasmid was modified to have the engineered NC-linkers from haXWnt8, haXWnt8c, and haXWnt8d synthesized by IDT, making hahWnt8, hahWnt8c, and hahWnt8d, respectively. The hahWnt8s-expressing cell lines were prepared and hLRP6_E1E2_ tetramer staining was confirmed as described above in comparison with the wild-type hWnt8, showing superior binding (Figure 1D and Figure 1—figure supplement 2).

The hLRP6_E1E2_ monomer binding was confirmed by incubating the hFzd5_CRD_-1TM cells that display Wnt8 variants with biotinylated hLRP6_E1E2_ at 30, 100, 300, 1000, 3000 nM concentrations, followed by washes with PFE and labeling with 50 nM SA647 (binding) and 1:50 FLAG-FITC (Wnt expression). The cells were again washed with PFE twice and analyzed by the flow cytometer. Two independent experiments were performed in triplicate, and the representative experiment was shown with Prism 9 (GraphPad).

### Preparation of recombinant XWnt8 and haXWnt8s

The five haXWnt8 variants (residues 22-338) were cloned into the pActinCD8 vector as previously described (Janda et al., 2012). *Drosophila* S2 cells were maintained in Schneider’s *Drosophila* Medium (Gibco) supplemented with 10% (v/v) FBS and GlutaMAX at 27°C with ambient CO_2_. The cells were co-transfected with wild-type or variants XWnt8, the pActinCD8 mFzd8_CRD_-Fc vector (Janda et al., 2012), and pCoBlast using the calcium phosphate precipitation kit (Invitrogen) according to the manufacturer’s protocol. Each cell line was bulk selected for three weeks with a gradual increase of Blasticidin (Invitrogen) concentrations; 5 μg/ml, 10 μg/ml, 20 μg/ml, for the first, second, and the third week, respectively. Expression levels were checked by western blotting using Rabbit *Xenopus laevis* Protein Wnt-8 Polyclonal Antibody (MyBioSource, MBS1497175) as a primary antibody and Dako Rabbit Anti-Goat Immunoglobulins/HRP (Agilent, P0449) as a secondary antibody, where haXWnt8 showed the best expression while haXWnt8d had substantial degradation. After the selection, the haXWnt8 cell line was transferred to conical flasks for gentle agitation, and when the culture volumes reached ∼50 ml, the cells were gradually expanded in shaker flasks with ESF 921 Insect Cell Culture Medium supplemented with 20 μg/ml Blasticidin to a final volume of 1L and incubated for 5 days.

XWnt8 and haXWnt8 were purified from clarified culture supernatants (Janda et al., 2017; Tsutsumi et al., 2020). The XWnt8-mFzd8_CRD_-Fc complex was captured by rProtein A Sepharose Fast Flow (Cytiva). After washing the sepharose resin with 10 column volumes HBS, XWnt8 was eluted with HBS containing 0.1% (v/v) n-dodecyl β-D-maltoside (DDM, Anatrace, Sol-Grade) and 500 mM NaCl, while the mFzd8_CRD_-Fc remained bound to the resin. Wnts were further purified by Con A agarose (Tsutsumi et al., 2020).

### TOPflash signaling assay

To confirm the affinity enhancing mutations do not interfere with XWnt8 activity, SuperTopFlash HEK293 (293STF) cells were stimulated with a gradient concentration of haXWnt8, wild-type XWnt8, and next-generation surrogate (NGS) Wnt in the presence or absence of R-spondin 2 (Janda et al., 2017; Miao et al., 2020). The agonist concentrations were 0.39 nM to 50 nM for XWnt8s and 0.016 nM to 2 nM for NGS Wnt, respectively, with two-fold serial dilution. 293STF were seeded into 96-well plates and maintained in DMEM media supplemented with 10% (v/v) FBS one day before Wnt stimulation. Cells were stimulated on the next day and lysed after 24 hrs following manufacturer’s protocol in Dual Luciferase Assay kit (Promega). Luminance signal readings were performed using or SpectraMax i3x (Molecular Devices). Two independent experiments were performed in triplicate, and the representative experiment was plotted and analyzed by Prism 9.

### Preparation of the ligand-receptor complex for structural study

The S2 cell line expressing hLRP6_E1E2_ was prepared as described above with the pActinCD8 and pCoBlast vectors. The S2 haXWnt8-mFz8_CRD_-Fc cells and hLRP6_E1E2_ cells were mixed at the cell ratio of 1:3, and co-cultured for 5 days. The complex was captured by rProtein A Sepharose, and eluted by adding in-house 3C protease, while the cleaved Fc remained bound to the resin and hLRP6_E1E2_ was affinity-purified by haXWnt8 bound to mFzd8_CRD_. The elution was concentrated down to 500 uL and injected to a Superdex 200 10/300 GL column equilibrated with HBS containing 2 mM CaCl_2_. The peak fraction with a retention volume corresponding to the 1:1:1 complex was concentrated to 0.5 mg/ml for cryo-EM analysis.

### Cryo-EM specimen preparation and data collection

3 µL sample was applied onto glow-discharged 300 mesh gold grids (UltrAuFoil R1.2/1.3). Excess sample was blotted to a filter paper for 3 sec with botting force of 3, before plunge-freezing with a Vitrobot Mark IV (Thermo Fisher Scientific) at 4°C and 100% humidity. The cryo-EM movies were collected using a Titan Krios (Thermo Fisher Scientific) operated at 300 kV equipped with Gatan K3 camera in counting mode. Nominal magnification was set to 29,000x, corresponding to a pixel size of 0.8521 Å and a calibrated magnification of 58,680x. Movies were recorded using SerialEM (Mastronarde, 2005) for 2.5 sec with 0.05 sec exposure per frame at an exposure rate of ∼16 electrons/pixel/sec at the specimen, and the nominal defocus range between -1.0 and -2.0 μm on gold support. A beam-image shift was used with an active calibration to collect 9 movies from 9 holes per stage shift and autofocus.

### Cryo-EM data analysis

Collected cryo-EM movies were processed and assessed with cryoSPARC Live. Patch motion correction was performed for the movies with a native pixel size of 0.8521 Å, and patch contrast transfer function (CTF) parameters were estimated with a default setting. 10,077 curated movies were used for the downstream processing by cryoSPARC (Punjani et al., 2017) (Figure 2—figure supplement 1B). Particles were picked with 2D templates from preliminary 2D classification and extracted with a box size of 352 pixels, binned to 132 pixels. Multiple rounds of 2D classifications, multi-class *ab intio* 3D reconstructions, and non-uniform 3D refinements yielded a preliminary 3D reconstruction with 43,337 particles. Using this particle set, three rounds of topaz (Bepler et al., 2019) picking and extraction were performed with a default setting using 918 micrographs and 50-200 expected particles per micrograph for training. This yielded a total of 3,840,364 particles including duplicated particles with the 352/132 binned box. After 2D classifications, multi-class *ab initio* 3D reconstruction and removing duplicates with 20 Å minimum separation distance, 276,237 particles were extracted with a box size of 352 pixels without binning. Further 2D classifications and *ab initio* 3D reconstructions identified 82,235 particles which were re-extracted with the 352-pixel box without binning, and beamtilt was fitted by Global CTF Refinement for each image shift group. Local 3D non-uniform refinement was performed using the mask covering the whole complex. The map was sharpened by deepEMhancer (Sanchez-Garcia et al., 2020) for visualization. The reported nominal resolution was 3.8 Å as determined by the 0.143 gold-standard FSC cutoff using 3DFSC (Zi Tan et al., 2017). As expected from the 2D class averages with preferred particle orientation (Figure 2B and Figure 2—figure supplement 1C), the map has a large variation in the directional resolutions, as well as highly diverged local resolutions, with the LRP6_E1E2_ density better defined (Figure 2—figure supplement 1D-F). The 3D map also indicates exposure to the air-water interface around XWnt8’s lipid (Figure 2—figure supplement 1E).

### Model building and refinement

The crystal structures of XWnt8-mFzd8_CRD_ (PDB 4F0A) and hLRP6_E1E2_ (PDB 3S94) were manually placed into the cryo-EM map using UCSF ChimeraX (Goddard et al., 2018). To facilitate map interpretation beyond the docking model, the missing N-terminal loop and NC-linker were modeled in the cryo-EM density using Coot (Emsley and Cowtan, 2004). Clearly distorted regions from the initial model, represented by the Wnt index finger, were flexibly fitted. We note that, based on the docked initial structures, the regions with weak cryo-EM density such as the Wnt thumb were kept in the final model but with lower occupancy (0.2). To avoid overinterpretation, all sidechains were truncated to Cβ except for prolines, cysteines and the palmitoleate (PAM)-modified serine. The PAM acyl torsion angles were also manually adjusted from the initial model. The model was refined with Phenix (Liebschner et al., 2019) using global minimization and B-factor with the starting model as a reference model. The blurred map with a 200 Å^2^ B-factor was used for the refinement to avoid overfitting, and the model resolution was estimated to be 9.7 Å with the model-FSC correlation cut-off of 0.5. AlphaFold2 (Jumper et al., 2021) was used to model hLRP6_E1E4_ for superimposition to the XWnt8-mFzd8_CRD_-hLRP6_E1E2_ model. The structure and map were visualized using UCSF ChimeraX and PyMol (Schrödinger, LLC). The secondary structures in the figures were assigned using DSSP (Joosten et al., 2011).

### Peptide synthesis

Peptide synthesis was outsourced to ChemParter (Shanghai, China); all the linker peptides were synthesized using standard 9-fluorenylmethoxycarbonyl protocol as described earlier (Stanger et al., 2012; Zhang et al., 2009). They were purified by reverse-phase HPLC. Peptide quality (>90% purity) was verified by liquid chromatography coupled to mass spectroscopy (LC-MS).

### Protein preparation for ELISA

Human LRP6 extracellular fragments, hLRP6_E1E2_ (residues 20-631) and hLRP6_E3E4_ (residues 631-1253), carrying a C-terminal hexahistidine-tag were secreted from *Trichoplusia ni* cells. Secreted proteins were isolated with an affinity column of immobilized anti-His-tag mAbs and eluted with buffer containing 50 mM sodium acetate pH 4.8 and 300 mM NaCl. The eluted proteins are further purified by SEC on a Superdex S200 column. The SEC column was equilibrated with buffer containing 10 mM sodium citrate pH 5.6 and 300 mM NaCl for hLRP6_E1E2_, and 10 mM sodium cacodylate pH 6.5 and 300 mM NaCl for hLRP6_E3E4_. Monomeric fractions were pooled, concentrated to ∼10 mg/ml, flash-frozen, and stored at -80°C until further usage.

LRP6-binding Fabs YW210 and YW211 were expressed as follows; 10 ml of inoculation culture was grown from *E. coli* 67A6 transformed with a Fab-encoding plasmid in Luria Broth (LB) media with 50 μg/ml carbenicillin overnight at 30°C. The inoculation culture was added to 500 ml of Soy CRAP media with 50 μg/ml carbenicillin and incubated overnight at 30°C with shaking. Expression cultures were spun down at 10,000 *g* for 10 min. Cell pellets were resuspended in buffer containing 200 ml PBS, 25 mM EDTA and Complete Protease Inhibitor Cocktail tablets (Roche, 1 tablet per 50 ml buffer). The mixture was homogenized and then passed twice through a microfluidizer. The suspension was centrifuged at 20,000 *g* for 45 min. The protein was loaded onto a Protein G column (Cytiva, 5 ml) equilibrated with PBS. The column was then washed with PBS and eluted with 0.6% (v/v) acetic acid. Proteins were subsequently purified by a HiTrap SP HP column (Cytiva) in buffer containing 25 mM MES pH 5.5 and eluted with a linear gradient of buffer containing 25 mM MES pH 5.5 and 500 mM NaCl. Monomeric fractions were pooled, concentrated to ∼150 μM, flash-frozen, and stored at -80°C until further usage. H07 VHH was provided by the Genentech Biomolecular Resources.

### ELISA

ELISA was carried out on NUNC Maxisorp plates coated with 5 μg/ml neutravidin. The blocking was performed with 0.5% (w/v) BSA in PBS for 2 hrs at room temperature, and 1 μg of biotinylated linker peptide per well was immobilized. Serial dilutions of hexahistidine-tagged LRP6 domains or Reck (R&D systems) were prepared in ELISA buffer (PBS, 0.5% (w/v) BSA, and 0.05% (v/v) tween 20) and added to the wells. The reaction mixture was incubated for 45 min at room temperature and washed with ELISA buffer. Horseradish peroxidase-conjugated anti-His antibody (Abcam, ab1187) was diluted in ELISA buffer at 1:5000 and added to the wells for 45 min incubation. Linker peptide-bound LRP6 domains or Reck were detected using TMB peroxidase substrate (Seracare, 2-component system), and the plates were read at 450 nm using SpectraMax M5e (Molecular Devices). For the competition assays with Fab YW210, Fab YW211 or H07 VHH, a constant concentration of LRP6 domains at 3x EC50 (Table 2) was used. A serial dilution of competitors was added, and the bound His-tagged LRP6 domains were detected as described.

### TOPbrite dual-luciferase Wnt reporter assays

HEK293 cells with stably integrated firefly-luciferase-based Wnt reporter (TOPbrite) (Zhang et al., 2009) and pRL-SV40 Renilla luciferase (Promega) were maintained in a 5% CO_2_ humidified incubator at 37°C in DMEM with nutrient mixture F12 (50:50), 10% (v/v) FBS, 2 mM GlutaMAX (Gibco) and 40 μg/ml hygromycin (Cellgro). Cells were grown for at least 24 hrs before any experiments. 20,000-40,000 cells/well in 50 μl medium were seeded in each well of clear-bottom white polystyrene 96-well plates (Falcon) and incubated for 24 hrs. Cells were then transfected with 0-25 ng of Wnt or Wnt-chimera-expressing constructs mixed with FuGENE HD in 10 μl OptiMEM, followed by incubation for 24-48 hrs. Treatment with peptides or LRP6 Fabs was done for 6 h before the assay measurement. Readout was obtained with 50 μl of Dual-Glo Luciferase Assay system (Promega) according to the manufacturer’s instructions on a Perkin Elmer EnVision multilabel reader. The ratios of firefly luminescence to Renilla luminescence were calculated, and in some instances normalized to control (non-treated) samples. Cell lines were tested for mycoplasma contamination and authenticated by single-nucleotide polymorphism (SNP) analysis.

### Western blot analysis

For Wnt expression assay, 4.0 × 10^6^ HEK293 luciferase reporter cells were seeded onto 10-cm dishes and transfected on the following day with 4 μg DNA encoding Wnt1, Wnt3a, Wnt5a, Wnt1_3a, Wnt3a_1 and Wnt5a_1 using FuGENE HD. Empty vector (pcDNA3.1) was used as control. After 48 hrs of incubation, culture medium was collected and cells were gently washed twice with 5 ml cold PBS and lysed in 0.5 ml of lysis buffer (PBS, 1% (v/v) Triton X-100 and protease inhibitor cocktail (Roche)). Collected lysate was centrifuged for 5 min at 18,400 *g* to remove cell debris, and supernatant was used to determine total protein concentration by BCA assay (Thermo Fisher Scientific). 10 μg of total protein was loaded onto an SDS-PAGE (4-12%) to assess Wnt protein levels. To evaluate secreted Wnt protein levels, the collected Wnt-conditioned medium (6 ml) was incubated with 50 μl Blue Sepharose beads (GE) on a rocking platform overnight at 4°C. The beads were washed three times with 500 μl PBS containing 0.1% (v/v) Tween 20 by centrifugation at 2,400 *g* for 5 min. 50 μl of 1x XT sample buffer including reducing reagent (Bio-rad) was added to the beads and the sample was boiled for 5 min at 95°C before loading onto an SDS-PAGE (4-12%). Proteins were then transferred to nitrocellulose membrane and probed with antibodies against Wnt1 (Abcam), Wnt3a (Cell Signaling), Wnt 5a/b (Cell signaling) and HSP90 (Cell Signaling).

For the measurement of LRP6/β-catenin levels, 2.0 × 10^5^ HEK293 luciferase reporter cells were seeded onto 24-well plates and transfected next day with 250 ng of DNA encoding Wnt proteins and chimeras as described above using FuGENE HD. After 24 hrs incubation in regular condition, cells were gently washed twice with 500 μl cold PBS and directly lysed in 50 μl of lysis buffer (PBS, 1% (v/v) Triton X-100 and protease inhibitor cocktail (Roche)). Collected lysate was centrifuged for 5 min at 18,400 *g* to remove cell debris and supernatant was used to determine total protein concentration by BCA assay. 15 μg of total protein was loaded onto an SDS-PAGE (4-12%) to assess LRP6 and β-catenin protein levels using phospho- and total LRP6 (Cell Signaling), active- and total β-catenin (Cell Signaling) and GAPDH (Cell Signaling) antibodies.

Antibodies were purchased from Abcam (Wnt1: ab15251 (Lot number GR3315807-3, dilution 1:1000)) and Cell Signaling Technology (Wnt3a: 2391S (Lot number 2, dilution 1:1000), Wnt5a/b (C27E8): 2530T (Lot number 4, dilution 1:1000), HSP90 (C45G5): 4877S (Lot number 5, dilution 1:2000), GAPDH (14C10): 2118S (Lot number 14, dilution 1:2000), LRP6 (C4C7): 2560S (Lot number 11, dilution 1:1000), Phospho-LRP6 (Ser1490): 2568S (Lot number 6, dilution 1:500), β-catenin (D10A8): 8480S (Lot number 5, 1:1000) and Non-phospho (active) β -catenin (Ser33/37/Thr41) (D13A1): 8814S (Lot number 7, 1:500).

### Quantitative RT-PCR analysis

4.0 × 10^5^ HEK293 cells were seeded onto 12-well plates and transfected next day with 500 ng of DNA encoding Wnt proteins and chimeras, Wnt1, Wnt3a, Wnt5a, Wnt1_3a, Wnt3a_1 and Wnt5a_1, using FuGENE HD. After 24 hrs incubation, cells were gently washed twice with 500 μl cold PBS and RNA was isolated using the RNeasy kit (Qiagen), as instructed by the manufacturer’s manual. Briefly, cells were lysed in RLT buffer including β-mercaptoethanol, which was followed by addition of 70% ethanol. The sample was then transferred to the column provided in the kit. After several washing steps, RNA was eluted using RNAse-free water. The real-time PCR reactions were performed with the TaqMan RNA-to-C_T_ 1-Step Kit (Applied Biosystems) by preparing samples as follows (total 10 μl reaction): 5.0 μl of 2X TaqMan RT-PCR mix, 0.5 μl of 20X TaqMan gene expression probe, 0.25 μl of 40X TaqMan RT enzyme mix, 50 ng of RNA (2 μl, 25 ng/μl), 2.25 μl of nuclease-free water. The reaction was initiated at 48 °C for reverse transcription for 15 min, followed by 45 amplification cycles (activation of AmpliTaq Gold DNA polymerase at 95 °C for 10 min, denaturation at 95 °C for 15 sec, and anneal/extend at 95 °C for 1 min). The assay was run on QuanStudio 7 Flex Real-Time PCR systems (Thermo Fisher Scientific). Relative RNA levels were calculated using the ΔΔC_*T*_ method and normalized to the level of housekeeping gene human HRPT1 within the same sample and further normalized to the sample from cells transfected with control empty vector. Taqman RNA-to CT 1 step kit (4392653) and gene expression probes (Axin2-FAM, Hs01063168-m1 and HPRT1-FAM, Hs02800695_m1) were purchased from Thermo Fisher Scientific.

## Material availability

The unique materials generated in this study will be available from the corresponding authors by request.

## Data availability

The cryo-EM map has been deposited in the Electron Microscopy Data Bank under accession code EMD-26989, and the model coordinate has been deposited to Protein Data Bank under accession number 8CTG.

## Acknowledgments

We thank members of the Garcia and Hannoush Labs for thoughtful discussion and helpful feedback. Cryo-EM data were collected at the Stanford cryo-EM center (cEMc). We thank Drs. Elizabeth Montabana, Yee-Ting Li, Aimin Song and Jeff Tom for their support. K.C.G. is an investigator of the Howard Hughes Medical Institute and the Younger Family Chair and is supported by the NIH funding (R01DK115728), the Mathers Foundation, and the Ludwig Institute.

## Competing interests

K.C.G. and C.Y.J. are co-founders of Surrozen, Inc.

## Author contributions

Conceptualization: N.T., S.H., S.H., C.Y.J., R.N.H., and K.C.G. Methodology: N.T., S.H., S.H., C.R.G., C.Y.J., R.N.H. and K.C.G. Investigation: N.T., S.H., S.H., D.W., N.W., Y.M., C.R.G., N.A.C., K.M.J., and C.Y.J. Funding acquisition: K.C.G. Project administration: R.N.H. and K.C.G. Supervision: R.N.H. and K.C.G. Writing – original draft: N.T., S.H., C.R.G., C.Y.J., R.N.H., and K.C.G. Writing – review and editing: all authors.

**Figure 1 —figure supplement 1.**
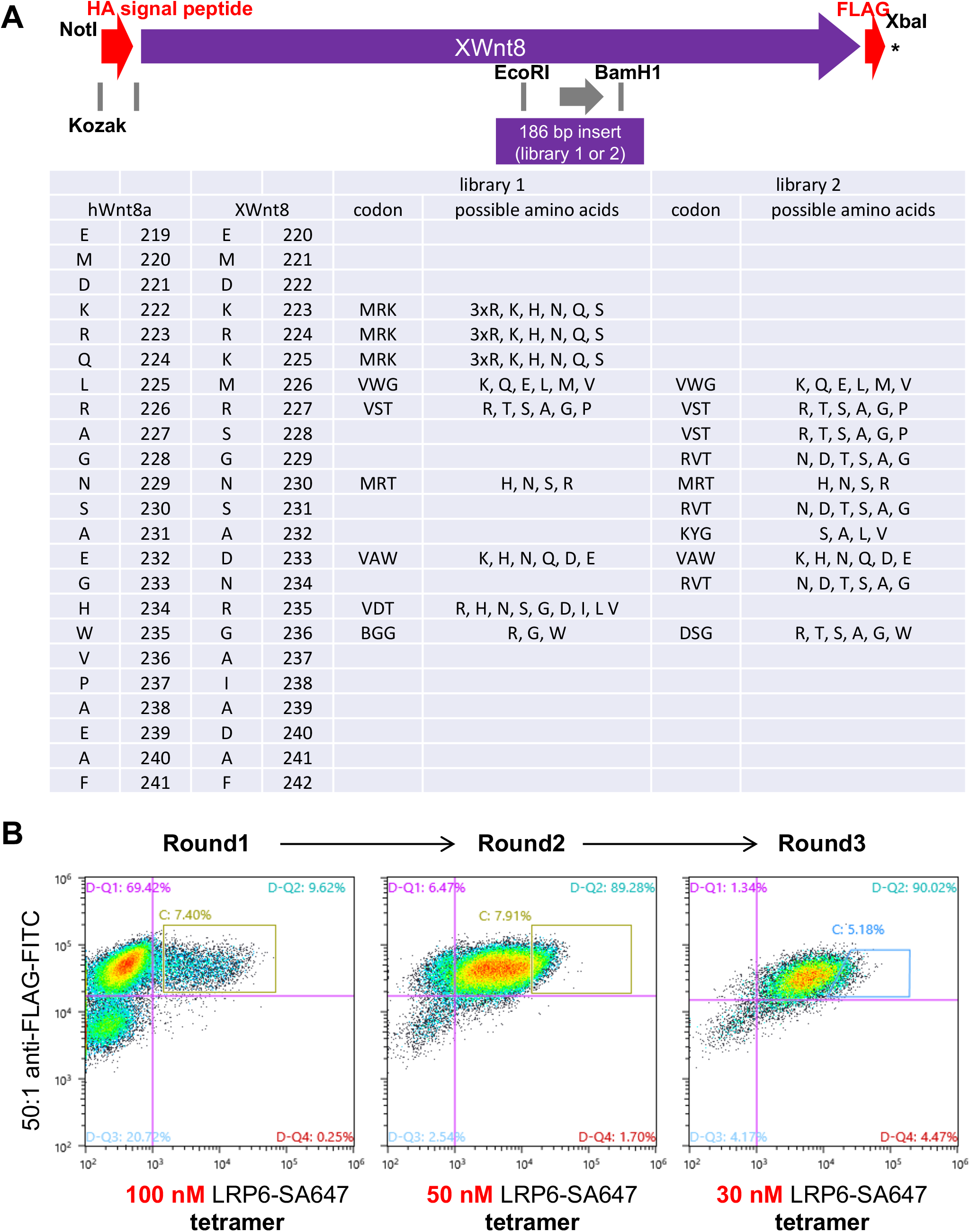
Construction and selection of the XWnt8 NC-linker library. (A) Design of the XWnt8 NC-linker libraries. (B) FACS plot and gating during the, round 1, 2, and 3 selections for the XWnt8 NC-linker library displayed on the mammalian cell surface.

**Figure 1—figure supplement 2.**
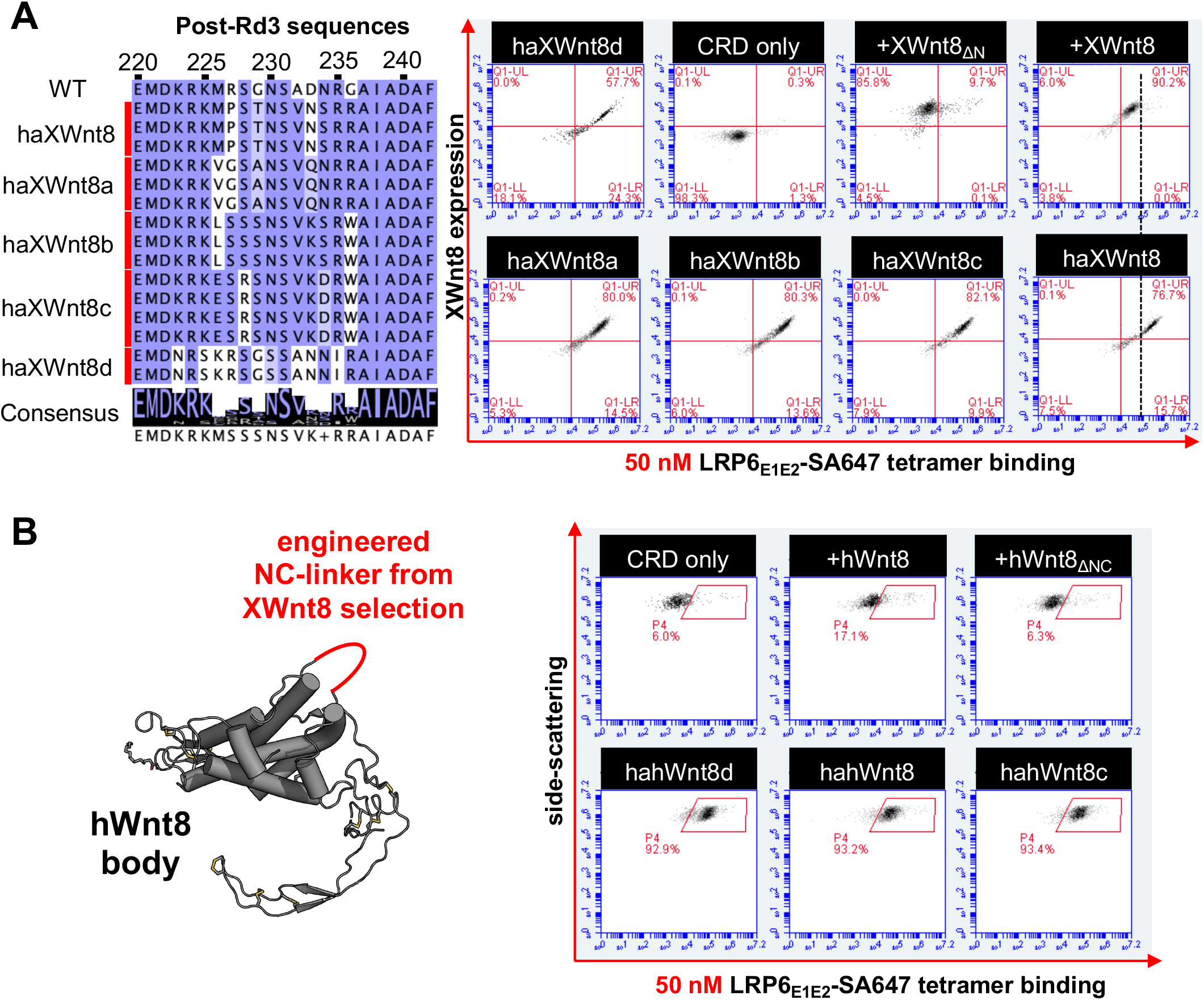
Single clones from the post-round 3 XWnt8 NC-linker library. (A) The NC-linker sequences for the five single clones after the round 3 selection in comparison with the wild-type sequence (left), and their hLRP6_E1E2_ tetramer binding on the engineered cells (right). (B) The schematic of the NC-linker grafting experiment from haXWnt8s to hWnt8 (left), and the chimeras’ hLRP6_E1E2_ tetramer binding on the engineered cells (right).

**Figure 2 —figure supplement 1.**
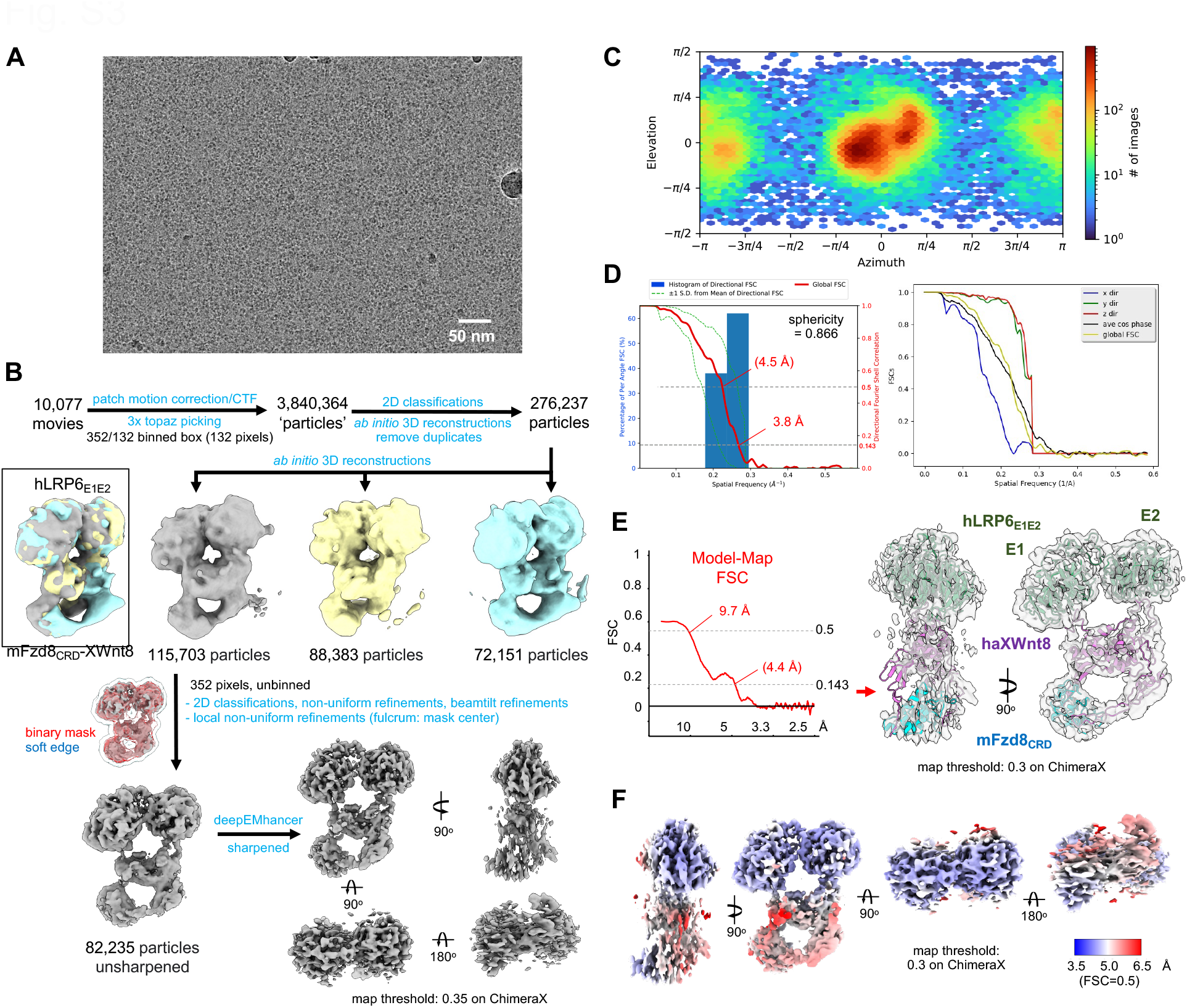
Cryo-EM data analysis. (A) Representative micrograph from cryo-EM data collection. (B) Cryo-EM data analysis workflow with representative 3D reconstructions during the data processing. (C) Angular distribution of the aligned particles in the final 3D reconstruction output from cryoSPARC. (D) Overall and directional gold-standard FSC curves calculated using 3DFSC. (E) The model-map FSC curve made with auto-masking by Phenix (left), and the cryo-EM map overlayed on the model (right). The red arrow on the right panel indicates the region with possible exposure to the air-water interface. (F) Local resolution estimates by Phenix colored on the surface representation of the deepEMhancer sharpened map. Blue: higher resolution, red: lower resolution.

**Figure 3 —figure supplement 1.**
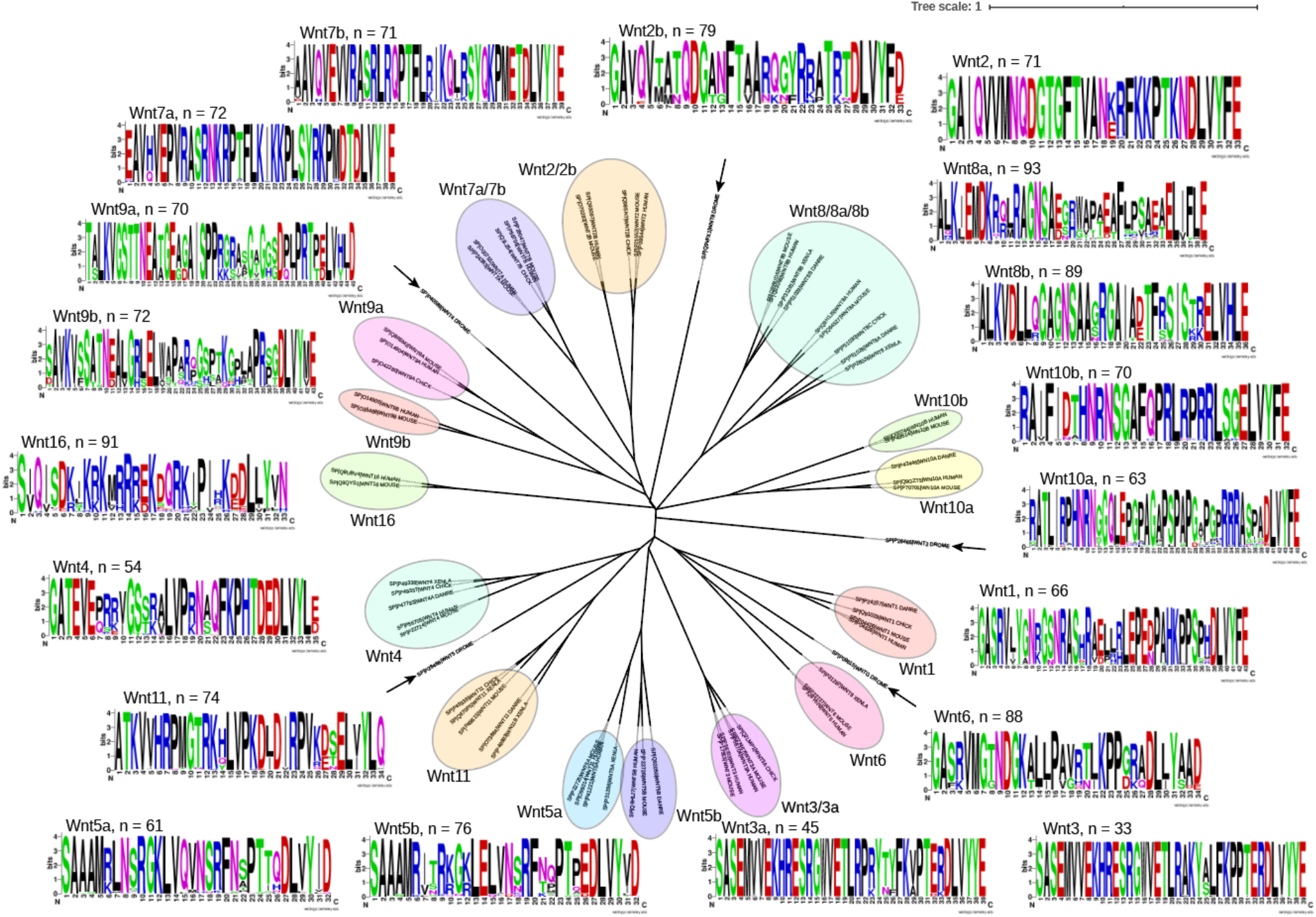
Phylogenetic analysis reveals evolutionary conservation of the NC-linker. The sequences of the NC-linker of Wnts from *Homo sapiens* (humans), *Mus musculus* (mouse), *Gallus gallus* (chicken), *Danio rerio* (Zebrafish), *Xenopus laevis* (African clawed frog) and *Drosophila melanogaster* (Fruit fly) were aligned and represented as phylogenetic tree. Arrows indicate *D. melanogaster* sequences. Weblogos are shown for the NC-linker sequence of each paralog from several species (number is indicated).

**Figure 3—figure supplement 2.**
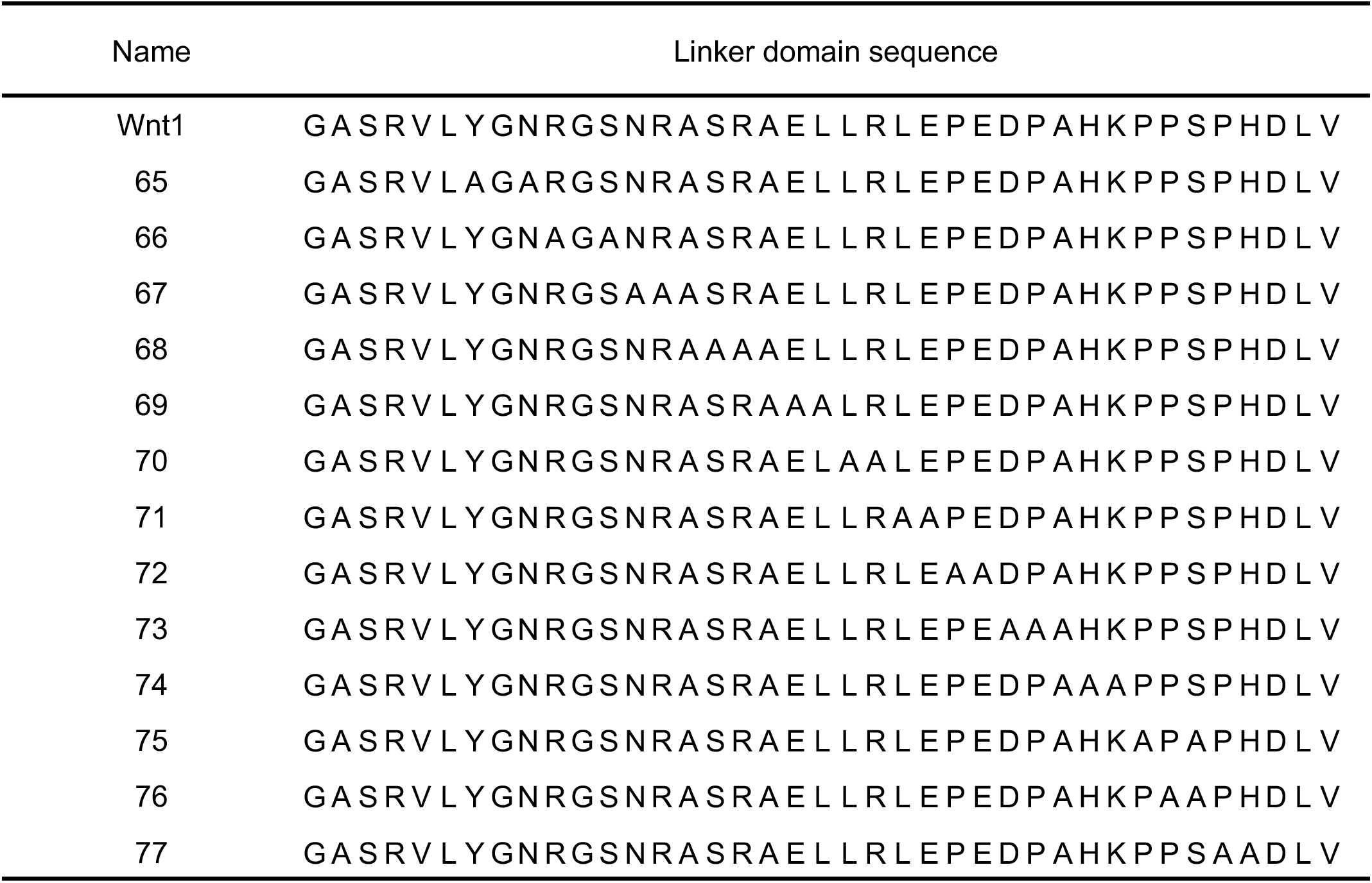
Sequences of the Ala mutants in Wnt1 NC-linker. Two residues were substituted for Ala in each mutant.

**Figure 3—figure supplement 3.**
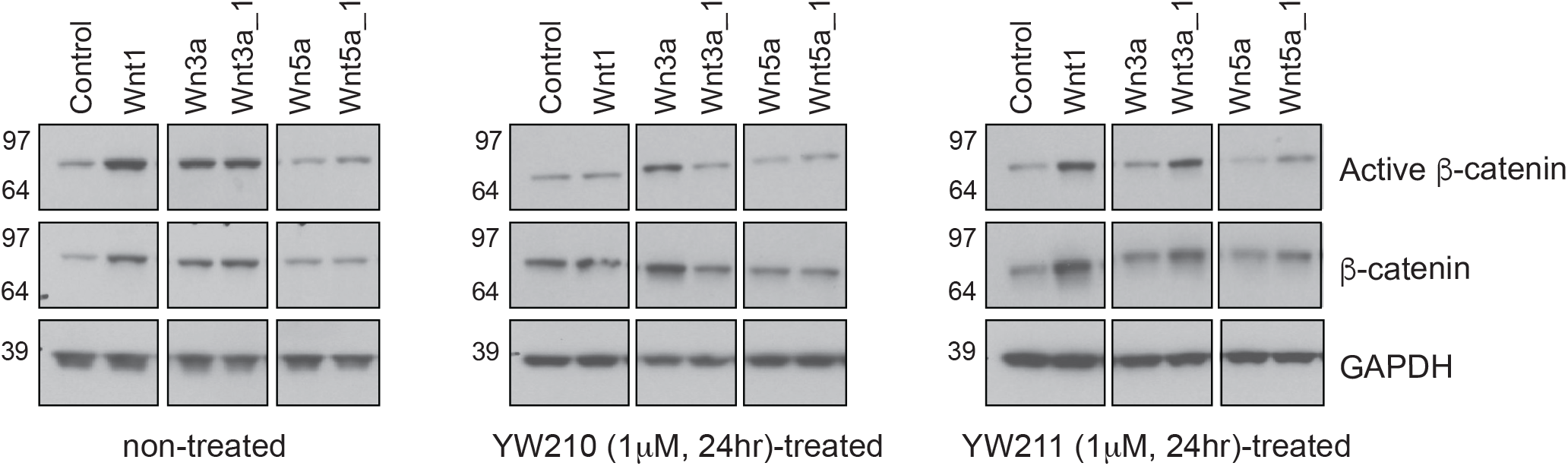
Representative Western blots showing b-catenin and LRP6 levels with overexpression of Wnts and Wnt chimeras in the presence of 1 μM Fab YW210 and Fab YW211. The treatment by FabYW210 significantly attenuated active b-catenin levels as compared to the control, whereas FabYW211 only affected Wnt3a signaling. Representative Western blot from one of three biological replicates. Figure 3—figure supplement 3—source data 1 | The raw western blot data showing β-catenin and LRP6 levels with overexpression of Wnts and Wnt chimeras in the presence of 1 μM Fab YW210 and Fab YW211. The boxes indicate the region shown in Figure 3—figure supplement 3.

**Figure 4 —figure supplement 1.**
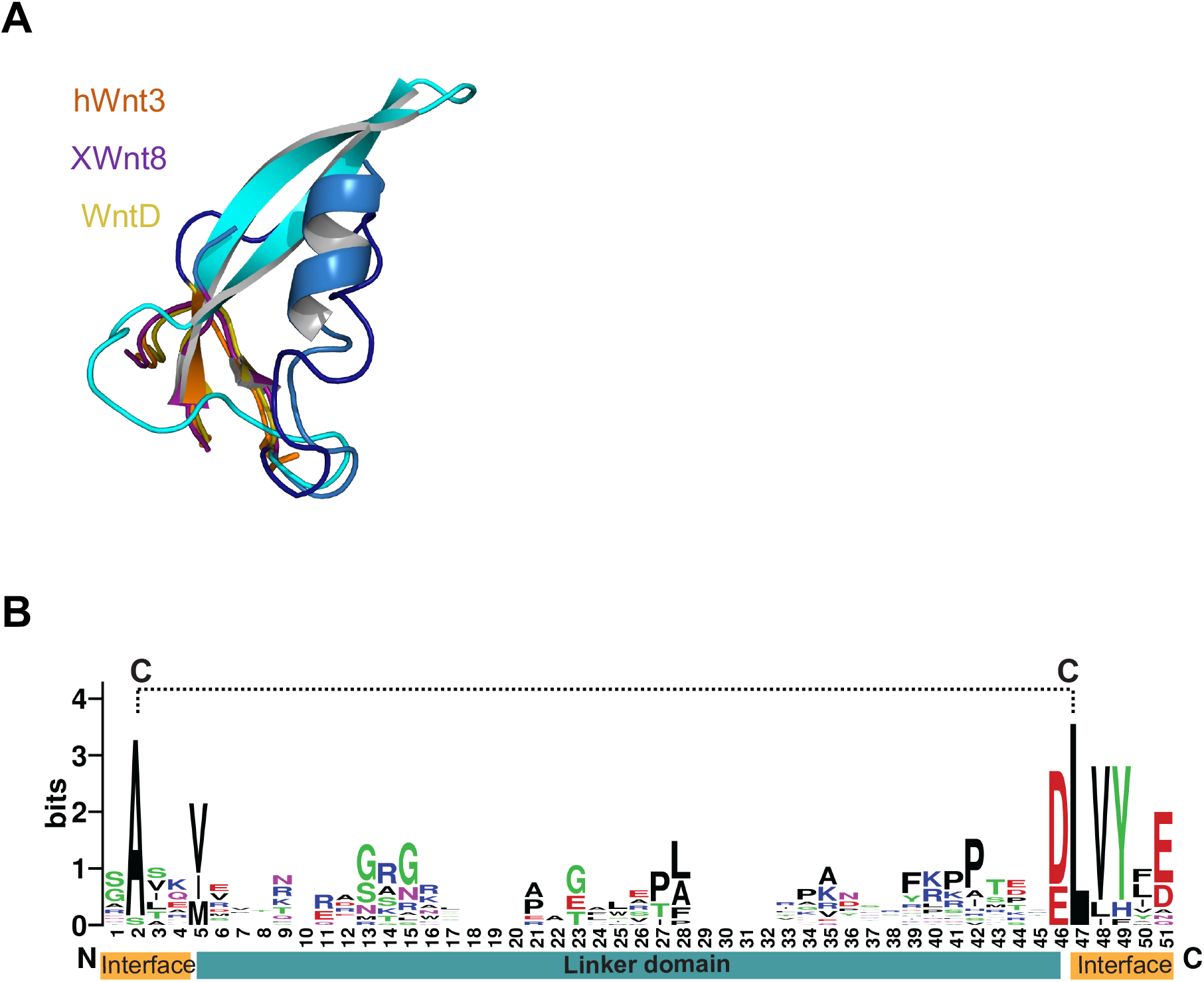
Sequence diversity of the NC-linker of Wnt isoforms. (A) Close-up view of the NC-linker. Superposition of all three available NC-linker in shades of blue and interface motifs in orange, purple and yellow (PDB hWnt3; 6AHY, XWnt8; 4F0A and WntD; 4KRR). (B) Weblogo shows the sequence conservation of the interface motifs and diversity of the NC-linker of all 19 human Wnts. Introduced disulfide bridge is indicated by black dashed line.

**Figure 4—figure supplement 2.**
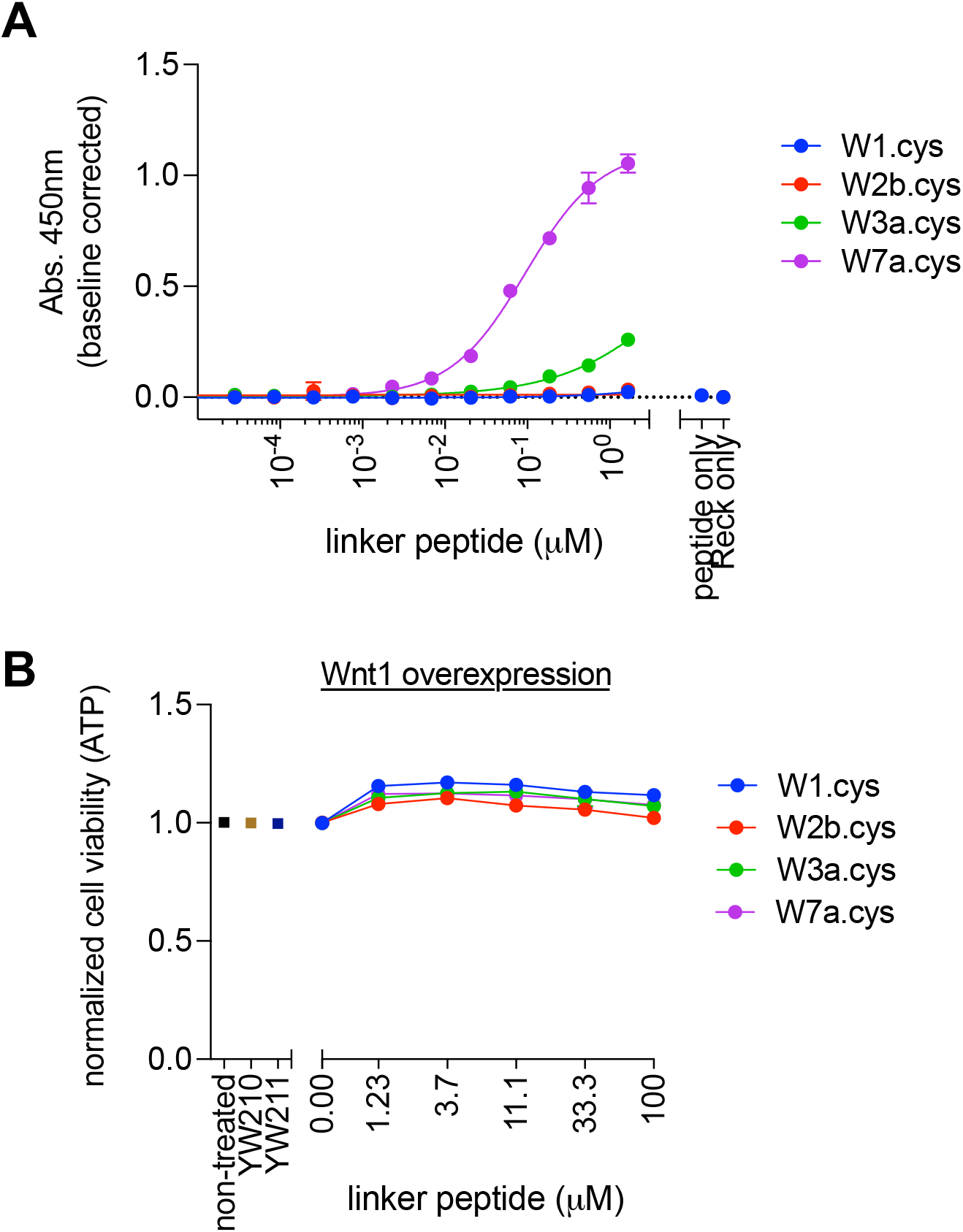
Specificity of W7a.cys peptide and effect of NC-linker peptides on cell viability. (A) ELISA showing the specific binding of W7a.cys peptide to co-receptor Reck. (B) The linker peptides did not cause toxicity to cells, indicating that the inhibitory effect of W1.cys peptide on Wnt1-mediated signaling (Figure 4D) was induced by the peptide. Bar and error bar represent the mean and SD of five technical replicates.

**Figure 5 —figure supplement 1.**
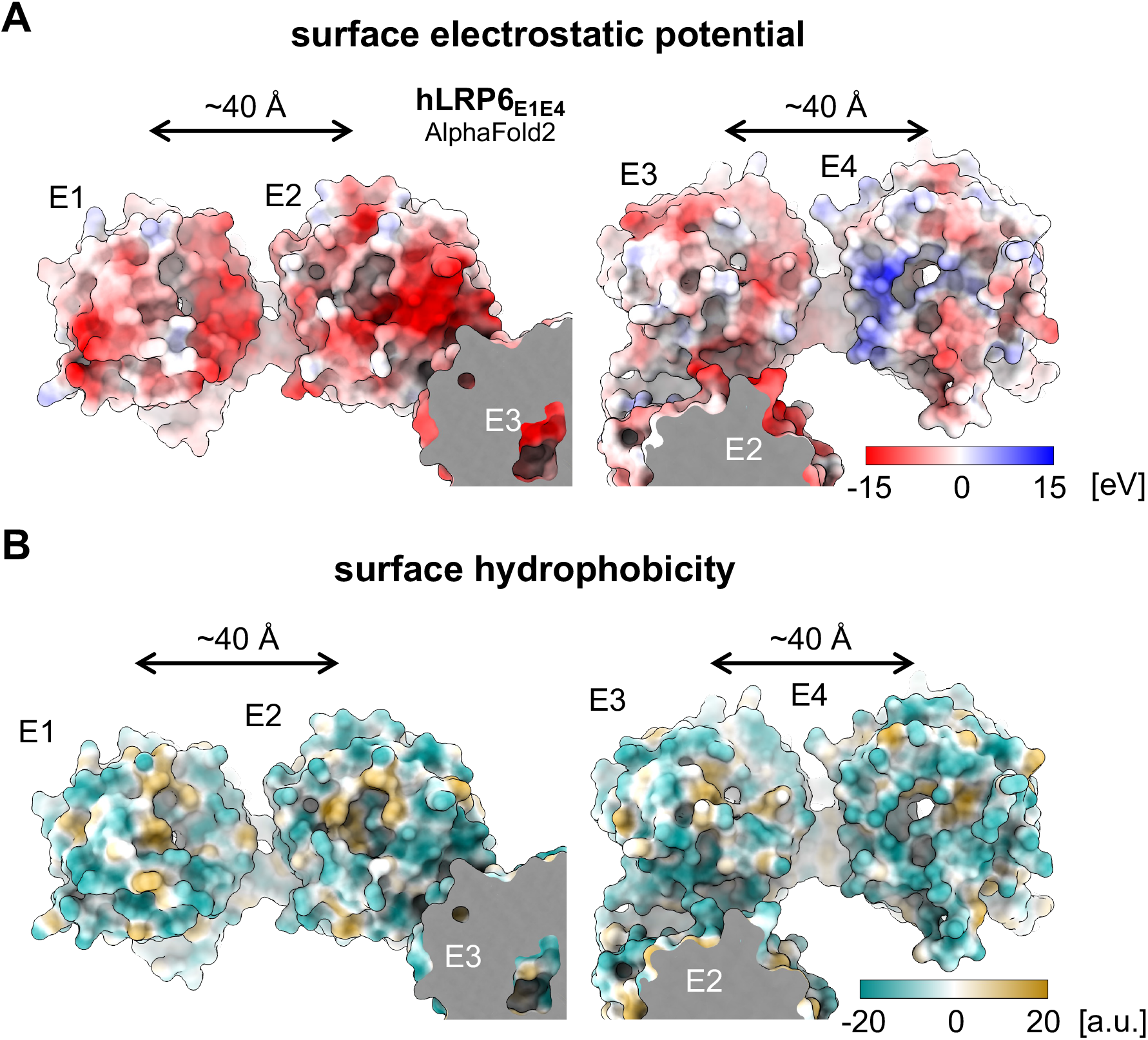
Modularity of LRP6_E1E2_ and LRP6_E3E4_. (A) Electrostatic potentials of hLRP6_E1E4_ looking down the β-propeller funnels. (B) Hydrophobicity of LRP6 funnels. The AlphaFold2 hLRP6_E1E4_ model in surface representation were colored as indicated in the panel, and the figure is prepared with UCSF ChimeraX. Red, more acidic; Blue, more basic; Brown, more hydrophobic; Green, more hydrophilic.

**Figure 5—figure supplement 2.**
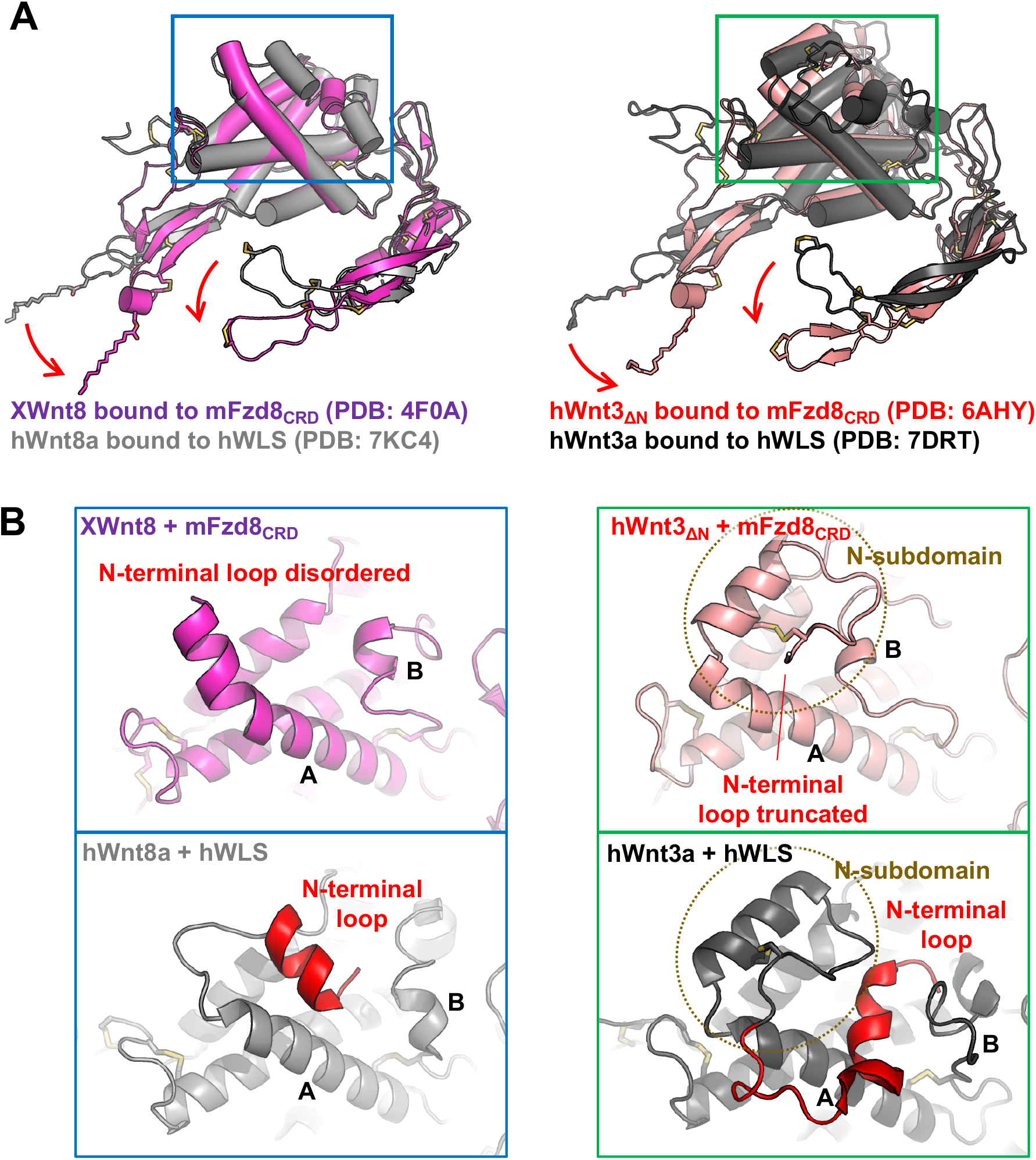
Comparison between the Wnt structures bound to Fzd or WLS. (A) Global structural rearrangement of the Wnts’ thumb and index fingers upon transfer from WLS to Fzd. (B) Freedom of Wnts’ N-terminal loops upon formation of the signaling-competent complex. The crystal structure of hWnt3-mFzd8_CRD_ was determined with N-terminal truncation of hWnt3 (Hirai et al., 2019). However, the binary complex was reported to be insoluble without this truncation, indicating that the hydrophobic N-terminal loop is structurally isolated from the Wnt core and exposed to the solvent when bound to mFzd8_CRD_.

